# Transient Acute Neuronal Activation Response Caused by High Concentrations of Oligonucleotides in the Cerebral Spinal Fluid

**DOI:** 10.1101/2025.02.13.638138

**Authors:** Mariana Bravo Hernandez, Curt Mazur, Hao Chen, Linda Fradkin, Justin Searcy, Sebastien Burel, Mackenzie Kelly, Dona Bruening, Jacqueline G. O’Rourke, Yuhang Cai, Jonathon Nguyen, Lisa Berman-Booty, Lendell Cummins, Hans Gaus, Berit Powers, Hien Zhao, Paymaan Jafar-Nejad, Scott Henry, Eric Swayze, Holly B. Kordasiewicz

## Abstract

Oligonucleotide (ON) therapeutics are promising as a disease-modifying therapy for central nervous system disorders. Intrathecal ON administration into the cerebral spinal fluid is a safe and effective delivery mode to the CNS. However, preclinical studies have shown acute toxicities following high-dose central ON delivery. Here we characterize a transient neurobehavioral change peaking 15 minutes after ON dosing and resolving after 120 minutes. Symptoms include shaking, muscle twitching, cramping, hyperactivity, stereotypic movements, hyperreactivity, vocalizations, tremors, convulsions, and seizures. These are collectively referred here as the acute neuronal activation response. Acute neuronal activation is observed in rats, mice, and non-human primates and is quantifiable using a simple scoring system. It is distinct from acute sedation seen with some phosphorothioate-modified antisense oligonucleotides, characterized by loss of spinal reflexes, ataxia, and sedation. The acute neuronal activation response is largely sequence-independent and is driven by ON chelation of divalent cations, particularly influenced by the divalent cations-to-ON ratio in the dosing solution. Acute neuronal activation can be safely mitigated by adjusting this ratio through magnesium supplementation in the ON formulation. We provide a comprehensive framework for quantifying and mitigating the acute neuronal activation response caused by high concentrations of centrally delivered ON therapeutics in preclinical species.

**GRAPHICAL ABSTRACT:** 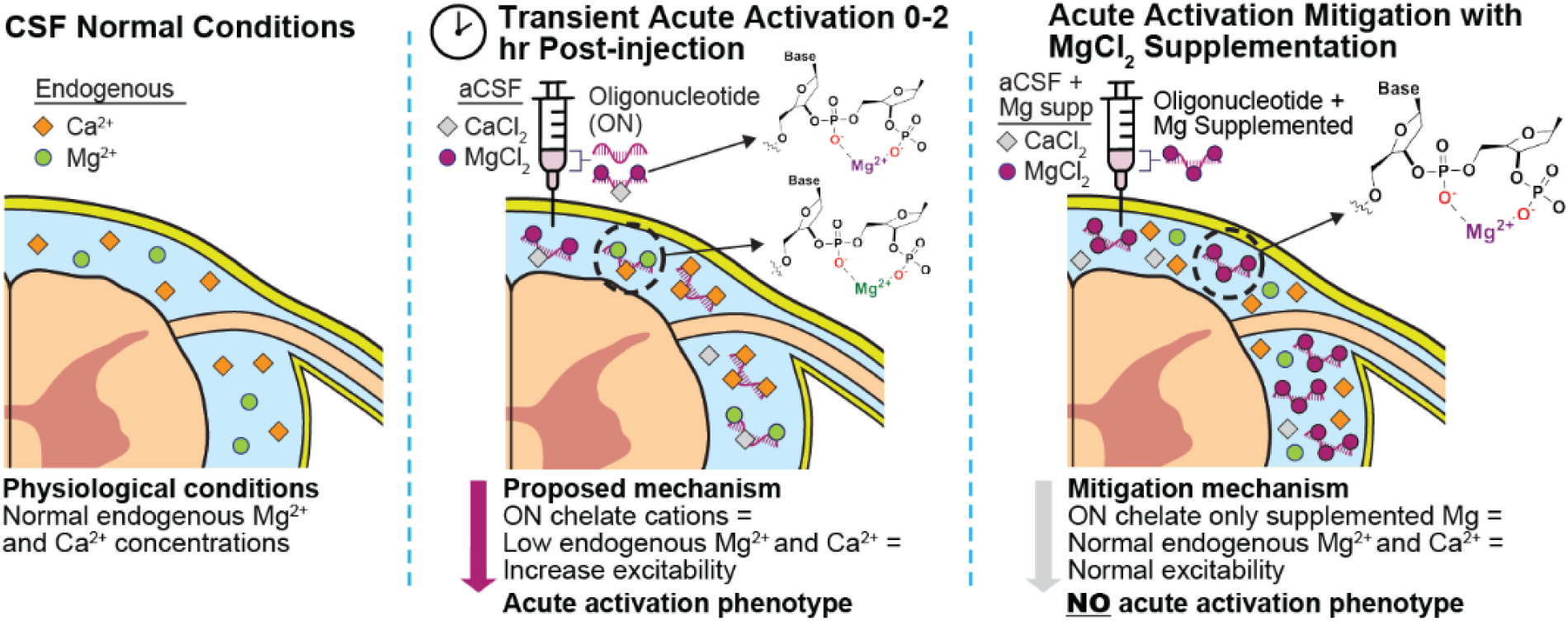

## INTRODUCTION

Oligonucleotide (ON) therapeutics hold tremendous promise as disease-modifying treatments for neurological diseases. Two single-stranded antisense oligonucleotides (ASOs) have been approved for central nervous system (CNS) indications (1–3), and multiple ASOs and double-stranded small interfering RNAs (siRNAs) are in clinical development for neurological diseases (4, 5). In people, ON therapeutics are directly delivered to the CSF through intrathecal (IT) delivery, enabling the targeting of RNAs throughout the CNS (6–8).

The local IT delivery of ON is advantageous because it circumvents the blood-brain barrier, enabling the distribution of highly charged ON macromolecules throughout the CNS (6–8). One consideration is that this delivery route leads to transient, high local concentrations of ON in the CSF and adjacent parenchyma in the minutes and hours immediately following dosing near the injection site. These high concentrations can lead to acute, transient behavioral responses. These transient responses have been reported as a mix of sedation-like behaviors which include ataxia, hyporeflexia, motor deficits, muscle weakness, lethargy, and slowed breathing, and neuronal activation-like behaviors such as hyperactivity, muscle spasticity, tremors, and seizures (9–12). Typically, these symptoms are grouped into a single response and scoring system and do not consider the varying presentations and temporal relationships (9, 11, 12). Studying ON-induced acute responses in this manner has limited our understanding of the underlying mechanisms.

To triangulate on a likely single mechanism, we focus on responses observed immediately after dosing and predicted to involve neuronal activation, including shaking, muscle twitching, cramping, hyperactivity, stereotypic movements, hyperreactivity, vocalizations, tremors, convulsions, and seizure. By focusing on acute neuronal activation after dosing, we are able to describe a unique set of signs with a distinct temporal course and mechanism. We demonstrate this is different from the highly sequence-dependent acute sedation response of some centrally delivered phosporothioate oligonucleotides, which is described in a companion publication (13).

The acute neuronal activation response has been observed following CNS delivery of high concentrations of ON in preclinical species, including mice, rats, and non-human primates, and to our knowledge, has not been observed in human patients. The dosing volume for safe intrathecal delivery via lumbar puncture is limited in each species (typically 5-10 mL in mice, < 0.1 mL in rats, 1.0-2.0 mL in NHP, and 15-20 mL in adult humans) (14). Consequently, when ON for nonclinical safety assessments are formulated in aCSF at a given dose, this results in a significantly higher concentration of ON in smaller preclinical species compared to the same dose formulated for human studies.

The symptoms of ON acute neuronal activation response in preclinical species are similar to those seen in hypocalcemia and hypomagnesemia in humans (15–19). Phosphodiester and phosphorothioate backbone linkages in ON are polyanionic in nature and are known to bind divalent cations (20–22). aCSF contains a fixed amount of divalent cations (2.2mM), and higher concentrations of ON will lead to lower divalent cation-to-ON rations. Others have reported some acute responses of centrally delivered ON can be mitigated with cations (12, 23). Taken together, we hypothesized that the acute neuronal activation response, observed in animals following high-concentration ON delivery into the CSF, is caused by ON-induced chelation of endogenous divalent cations.

Here, we provide a comprehensive characterization of the acute neuronal activation response across rats, mice, and NHP. We developed a reliable, and species-specific scoring system to isolate and evaluate this activation phenotype. Additionally, we demonstrate that the acute neuronal activation response can be attributed to the chelation of endogenous cations associated with high ON drug concentrations. This response can be mitigated by supplementing the formulation with additional cations to buffer the ON chelating capacity, thereby minimizing its ability to sequester endogenous cations by improving the divalent cation-to-ASO ratio. Our findings demonstrate that the degree of ON saturation with divalent cations can be empirically defined, offering a practical approach to reducing the acute neuronal activation response in animals. The data described here are fundamental for translating findings and developing mitigation strategies across species.

## MATERIALS AND METHODS

### Oligonucleotides

Lyophilized ASOs A-Q or siRNA A (**Supplementary Table 1**) were dissolved in DPBS (catalog # 14190144, Gibco™), standard artificial cerebrospinal fluid (aCSF), or sodium-deficient artificial cerebrospinal fluid (dACSF) or modified artificial cerebrospinal fluid (mACSF) with or without different divalent cations-to-ASO ratios by adding MgCl_2_, or CaCl_2_ or MgCl_2_+CaCl_2_. Depending on the ASO and specific experiment, the final ASO concentrations ranged between 1 mg/mL (0.14 mM) to 100 mg/mL (14.1 mM) (**Supplementary Tables 3-11**). ASO dose solutions were formulated according to purity and sterilized by passage through a 0.2 um filter before dosing.

### Artificial cerebrospinal fluid formulations

#### Standard artificial cerebrospinal fluid (aCSF)

Stock aCSF was formulated containing: 0.32mM NaH_2_PO4·2H_2_O, 0.68 mM Na_2_HPO_4_, 150 mM NaCl, 3 mM KCl, 1.4 mM CaCl_2_·2H_2_O, 0.8 mM MgCl_2_·6H_2_O

#### Sodium deficient artificial cerebrospinal fluid (dACSF)

Stock dACSF was formulated containing: 0.32 mM NaH_2_PO4·2H_2_O, 0.68 mM Na_2_HPO4, 3 mM KCl, 1.4 mM CaCl_2_·2H_2_O 0.8mM MgCl_2_·6H2O with a variable amount of NaCl, 96 mM, 42 mM or 0 mM to control the solution tonicity after adding 40 mM MgCl_2_ alone, 14.1 mM ASO alone or 40 mM MgCl_2_ plus 14.1 mM ASO as noted in the specific experiment.

#### Modified artificial cerebrospinal fluid (mACSF)

Stock mACSF was formulated containing: 0.32 mM NaH_2_PO4·2H_2_O, 0.68 mM Na_2_HPO4, 3 mM KCl, 150 mM NaCl without MgCl_2_ or CaCl_2_.

#### Divalent Cations supplementation

MgCl_2_ and CaCl_2_ stock solutions were prepared individually in WFI water at a concentration of 2.5 M. To prepare the aCSF, dACSF, or mACSF containing the different divalent cations concentrations necessary for each experiment, a spike of MgCl_2_, CaCl_2_, or CaCl_2_ + MgCl_2_ (maintaining their physiological ratio Ca: Mg of 1.75:1) from the corresponding stock was done in the corresponding artificial cerebrospinal fluid. The final solutions were then used to dissolve the lyophilized ASO as indicated above.

### Animals

All rodent animal experiments were performed under a protocol approved by the Institutional Animal Care and Use Committee of Ionis Pharmaceuticals (Protocol # 2021-1164, and # 2021-1176). Female and male Sprague-Dawley rats with body weights of ∼200 - 230 and ∼260 - 380 g, respectively were obtained from Charles River laboratories (Sprague Dawley, strain code 687). Female and male C57BL/6 mice with body weights between 17-20 g were obtained from Taconic Bioscience (C57BL/6 B6, model B6-M).

Non-human primate experiments were performed at the Korea Institute of Toxicology (KIT) and were approved by their Institutional Animal Care and Use Committee. Eight female and eight male cynomolgus macaques of Vietnamese origin, with body weights of 2.6 ± 0.18 and 3.1 ± 0.59 kg respectively, were used, with approximate ages between 2-4 years old.

### Intrathecal (IT) delivery in rat

Catheterization of the lumbar intrathecal space was performed as described in *Mazur et al.* (24). Thirty or one hundred microliters of the corresponding ASO formulation or vehicle followed by 40 µl of the same vehicle were injected into the subarachnoid intrathecal space via the catheter over approximately 30 seconds. The specific volume, ASO formulation, and vehicle are described in each experiment and in the supplementary tables. Immediately after dosing, the animals were housed in their home cages to allow recovery from anesthesia.

### Intrathecal (IT) delivery in NHP

Cynomolgus monkeys were anesthetized with a ketamine and medetomidine cocktail and a reversal agent, atipamezole, was given following the completion of dosing. The corresponding ASO formulation or vehicle was administered via percutaneous intrathecal injection using a spinal needle at the lumbar level (target L5/L6 space) in the lateral recumbency position, as described before (6). The dose volume was fixed for all injections at 1 ml and administered over 1 min as a slow bolus.

### Intracerebroventricular (ICV) delivery mouse

Intracerebroventricular ASO delivery was performed as previously published (6). Briefly, Mice were placed in a stereotaxic frame and anesthetized with 2% isoflurane. The scalp and neck dorsal area were then shaved and disinfected. A small incision (approximately 1 cm) was made in the scalp, and the subcutaneous tissue and periosteum were scraped from the skull with a sterile cotton-tipped applicator. A 10 µl Hamilton micro syringe with a 26 G needle was driven through the skull at 0.3 mm anterior, 1.0 mm lateral to bregma, and 3.0 mm dorsoventral. Ten microliters of corresponding ASO solution or vehicle were injected a single time into the right lateral ventricle at a rate of 1 µl/s. After 3 min, the needle was slowly withdrawn, and the scalp incision was sutured. The mice were then allowed to recover from the anesthesia in their home cage.

### Acute Neuronal Activation Score evaluation

#### Rats

The acute neuronal activation scoring system in rats was designed to capture and group the signs and behaviors in levels based on their severity. We created 8 levels, from 0 to 7. The summary of the scoring system is described in **Table 1**. Briefly, Level 0 is an animal displaying no abnormal clinical signs or behaviors (alert or at rest performing maintenance behaviors). Level 1 is an animal that displays any of the following: a hunched back with signs of pain (pain defined by (25)), light shivering, spontaneous hindlimb twitching, hopping in place, and hind limb hypermotility while remaining in place. Level 2 would be assigned when the animal shows at least one of the following: a hunch-back while standing on its toes, severe shivering/shaking, hopping in place with mild vocalizations (low-pitched squeaking), or hypermotility in all limbs while remaining in place. For a level 3 rating, an animal must show: standing up in visible distress or intense hopping around the cage with mild vocalizing or all-limb hypermotility moving around the cage. For a level 4 score, an animal must be hyper-reactive to stimulus, such as cage mate movements and sounds, or display defensive/aggressive behavior and mild vocalizations. Animals with severe vocalizations (high-pitched, long, unmodulated vocal emissions) were scored as 5. A 6 was assigned when an animal had any kind of tremor or seizure, and a 7 was given if the animal died within the first two hours after dosing.

**Table 1.** Acute neuronal activation (aA) score system in rats after IT administration (0-7)

| aA score | Observable signs |
| --- | --- |
| 0 | No remarkable findings |
| 1 | Hunched in pain, shivering, spontaneous hindlimb twitches, hopping while remaining in place, hind limb hypermotility while remaining in place |
| 2 | Hunched and standing on toes, shivering/shaking, hopping while remaining in place and mild vocalizations (low-pitched squeaking), all limb hypermotility while remaining in place |
| 3 | Standing up in visible distress, intense hopping and mild vocalizations (low-pitched squeaking) moving around the cage, all limb hypermotility moving around the cage |
| 4 | Hyperreactivity to stimulus (cage mate movements, sounds, etc.), defensive behavior and mild vocalizations (low-pitched squeaking) |
| 5 | Severe vocalizations (high-pitched, long, unmodulated vocal emissions) |
| 6 | Any type of tremors or seizure |
| 7 | Death |

Rats are observed for 2 h after dosing, and the maximum score observed is recorded at 15, 30, 45, 60, 90, and 120 min after dosing. A higher score equates to a more severe response. The animals are scored always compared to the behavior of the control (vehicle) group. Pain was defined by the Rat Grimace scale (25).

#### Mice

Previously neurotoxicity score systems have been published in mice: The EvADINT system which scores together sedation and activation-like responses (12) and a modification of it for a seizure-like phenotype scoring (23). In the present study, we designed a scoring system to capture activation separately from sedation. Our scoring system is similar to that described for rats above in that the groupings and severity of the activation signs are assigned a single number from 0 to 7. This scoring system allowed us to evaluate the responses of mice at different time points in an accurate and simplified way. When a spectrum of clinical signs was observed, the most severe (highest score) was used for that subject. A 0 represents animals alert but at rest or performing maintenance activities such as sleeping, eating, grooming, or nesting. A score of 1 was assigned if animals had hunched backs with signs of pain (pain defined as(26)) or were freezing in the cage. A score of 2 was assigned when shivering or spontaneous shaking occurred. A score of 3 represented animals with stiff and raised tails and/or hyperactive scuttling around the cage. A score of 4 was used when severe limb stiffness that impaired forward movement was present (stiff limbs when walking). A score of 5 was given if they presented evident popcorning (vigorous jumping and hyperexcitability) (27) or hopping behavior. A 6 represented any kind of seizure. A score of 7 was given if the animal died during the 2-hour evaluation **Table 2**.

**Table 2.** Acute neuronal activation (aA) score system in mice after ICV administration (0-7)

| aA score | Observable signs |
| --- | --- |
| 0 | Alert at rest or performing maintenance activities (sleeping/eating/grooming/ nesting) |
| 1 | Hunched in pain/ freeze |
| 2 | Shivering or spontaneous shaking |
| 3 | Stiff and raised tail and/or hyperactive scuttling around the cage |
| 4 | Severe limb stiffness than impaired forward movement |
| 5 | Hopping or Popcorning (vigorous jumping and hyperexcitability) |
| 6 | Any type of seizures |
| 7 | Death |

Mice are observed for 2 hours after dosing, and the maximum score is recorded at 15, 30, 45, 60, 90, and 120 min. A higher score equates to a more severe response. Pain was defined by the Mouse Grimace scale (26)

#### Non-Human Primates

The acute neuronal activation scoring system for non-human primates (NHP) was similarly designed to capture and group the signs and behaviors in scores or levels based on their severity. We created 8 levels, from 0 to 7. When a spectrum of clinical signs were observed, the most severe (highest score) was used for that subject. A 0 represented no remarkable findings. To score a level 1, the NHP should show any of the following: tremor, twitch, salivation, self-biting, and/or urinary incontinence. Level 2 was given when the animals showed a combination of two signs of level 1 or when they presented with nystagmus alone. Level 3 was scored when tonic or clonic convulsions were present. If an NHP showed partial tonic-clonic convulsions a score of 4 was given. The presence of seizures scored a 5. The progression to severe seizures was scored a 6, and a 7 was given if the animal would require euthanasia. A summary of these descriptions is captured in **Table 3**.

**Table 3.** Acute neuronal activation (aA) score system in non-human Primates after lumbar puncture (0-7)

| **aA score** | **Observable signs** |
| --- | --- |
| 0 | No remarkable findings |
| 1 | Tremor, twitch, salivation, self biting, or urinary incontinence |
| 2 | Combination of two signs of the score one, or nystagmus |
| 3 | Tonic, or <u>clonic</u> convulsion |
| 4 | Tonic-<u>clonic</u> convulsion |
| 5 | Seizure |
| 6 | Severe seizures |
| 7 | Euthanasia required for human reasons |
| A higher score equates to a more severe response |  |

### Tissue collection

Mice and rats were euthanized at either 2 weeks or 8 weeks after dosing as described in the corresponding experiment. Brains were bisected sagittally to separate the hemispheres. A ∼1 mm wide piece of the parietal cortex was dissected from the right hemisphere, and a 3 mm length segment of the spinal cord (thoracic cord) was also dissected. Then, these tissue samples were frozen in dry ice and used to measure ASO concentration and mRNA expression of target or tolerability markers. The left hemisphere of the brain and another 3 mm segment of the spinal cord (thoracic) were immersion fixed in 10% neutral buffered formalin solution for histological processing and IHC staining.

### RT-qPCR

RNA was extracted using TRIzol® (TRIzol Reagent, Invitrogen, cat # 15596026). Then, total RNA was isolated using the RNeasy 96 Kit (Qiagen, Germantown, MD). RT-qPCR was performed using the EXPRESS One-Step Super-Script qRT-PCR kit. Gene-specific primers were designed (**Supplementary Table 12**) and purchased from IDT technologies (Coralville, IA). For ASO-treated animals, the expression level of *Malat1*, *Gfap*, *Aif1*, and *ATXN3* gene was normalized against *Gapdh* and this was further normalized to the mean level in vehicle-treated animals.

### Quantification of ASO tissue concentration

ASO tissue concentration was quantified and analyzed in rat CNS tissue as previously described (28–30). Samples were weighed, minced, and homogenized in a 96-well format plate. A calibration curve and quality control samples of known ASO concentrations were prepared and analyzed on the same plate. The ASO was extracted via a liquid-liquid extraction, and the aqueous layer was further processed. The aqueous layer was processed via solid phase extraction with a Strata X plate. Eluates were passed through a protein precipitation plate and dried out with nitrogen at 50◦C. Dried samples were reconstituted in 100 µl water containing 100 µM EDTA. The ASO concentration was quantified on an Agilent 6495 triple quadrupole LCMS.

### Immunohistochemistry

The left hemisphere and the thoracic spinal cord (3 mm length) were immersion fixed in 10% Neutral Buffered Formalin (Statlab, 28600-1) for 72 h, then, transferred to 70% ethanol for 48 h before processing. The tissue was processed into Paraplast Plus paraffin (Leica, 39602004) overnight on a Sakura Tissue-Tek tissue processor. After paraffin embedding, slides were cut at 5 μm, air dried overnight, and then dried at 60°C for 1 hour. ASO immunostaining was performed as previously reported (6, 8). Briefly, deparaffinization and hydration steps were performed on the slides before staining. The Thermo/Labvision Auto-stainer was used for antibody staining. The primary antibody used for ASO detection, a polyclonal rabbit anti-ASO antibody, was incubated for 1h using a 1:10 000 dilution (31). A donkey anti-rabbit HRP secondary was incubated (Jackson, 711-036-152) for 30 min followed by a DAB+ Substrate Chromogen System (Dako, K3468) for visualization. Finally, slides were counterstained with Hematoxylin for 30 s then dehydrated, cleared, and cover slipped with Micromount (Leica,3801731).

### Determination of ON binding to divalent cations

Binding of magnesium and calcium was assessed in a dialysis experiment utilizing thermo micro-dialysis tubes with a molecular weight cut-off of 3kDa **(Figure 8A)**. The utilized buffer systems were 10 mM sodium acetate buffer pH 7.2, and 150 mM NaCl. The ASO at a concentration of 10 mM was dialyzed in buffer solutions with increasing magnesium or calcium concentrations (0, 2, 5, 10, 20, 40, 80, 200 mM) overnight. The metal ion concentration before and after dialysis was determined using HILIC chromatography utilizing an evaporative light scattering detector (Agilent application note). Bound metal ion was plotted against initial metal ion concentration. Binding saturation ion concentrations were calculated using GraphPad Prism using one-site specific binding analysis.

### Data analysis and statistics

Data analysis was expressed as a mean ± s.d. GraphPad Prism version 10.0.0 for Windows, GraphPad Software, Boston, Massachusetts USA, or JMP®, Version 16. SAS Institute Inc., Cary, NC, 1989–2023 were used to plot the data and perform statistical analysis. The mean values from behavioral scores from individual groups were compared and analyzed using a non-parametric Mann-Whitney test or Kruskal-Wallis test. For other analyses a parametric Unpaired t-test, a one-way analysis of variance (ANOVA), or repeated measurements two-way ANOVA with a Tukey’s post hoc test. The corresponding analysis is described in the figure legends. The P values were adjusted for multiple comparisons, depending on the total number of treatment groups. The AUC for the time courses was calculated using the trapezoidal rule. ED50 and EC50 were calculated after fitting the data using nonlinear regression with the normalized response and variable slope using the Motulsky: Agonist vs. response -- Variable slope (four parameters) built on GraphPad Prism software and the F-test was used to compare the dose-response between groups. Significance was noted in each figure and figure legend when P≤ 0.05.

## RESULTS

### An immediate, transient, and body-weight-dependent acute neuronal activation response occurs following intrathecal oligonucleotide administration in rats

In rats, where high doses of ONs were delivered by IT bolus injection, muscle cramping, twitching, and vocalization have been observed immediately after dosing in some animals. The behavioral response was consistent with activation of the central nervous system and resolved quickly with no sequala. We termed this the acute neuronal activation response. To enable a precise analysis of the acute neuronal activation response, we devised a scoring system that captures the behaviors associated with the acute neuronal activation response that occurs immediately after IT dosing of ON in rats (Acute Activation Score [aA score], **Table 1**). To characterize this response, we administered to rats a very high intrathecal dose (3 mg dose, human equivalent dose of 1500 mg) of a single stranded ASO (ASO A, **Supplementary Table 1**), and scored their behavior in 15-minute intervals. The onset of acute neuronal activation response occurred immediately upon recovery from anesthesia. A clear temporal relationship of various signs and behaviors followed, generally declining over time, and resolving after 120 minutes (scores were binned in 3-time blocks; 0-15 minutes when peak responses were observed, 30-60 minutes, and 90-120 min, **Figure 1A**). Going forward, we will focus on the peak 15-minute timepoint and 3 mg dose of ON (100 mg/ml, 14.1 mM) unless otherwise noted.

**Figure 1.**
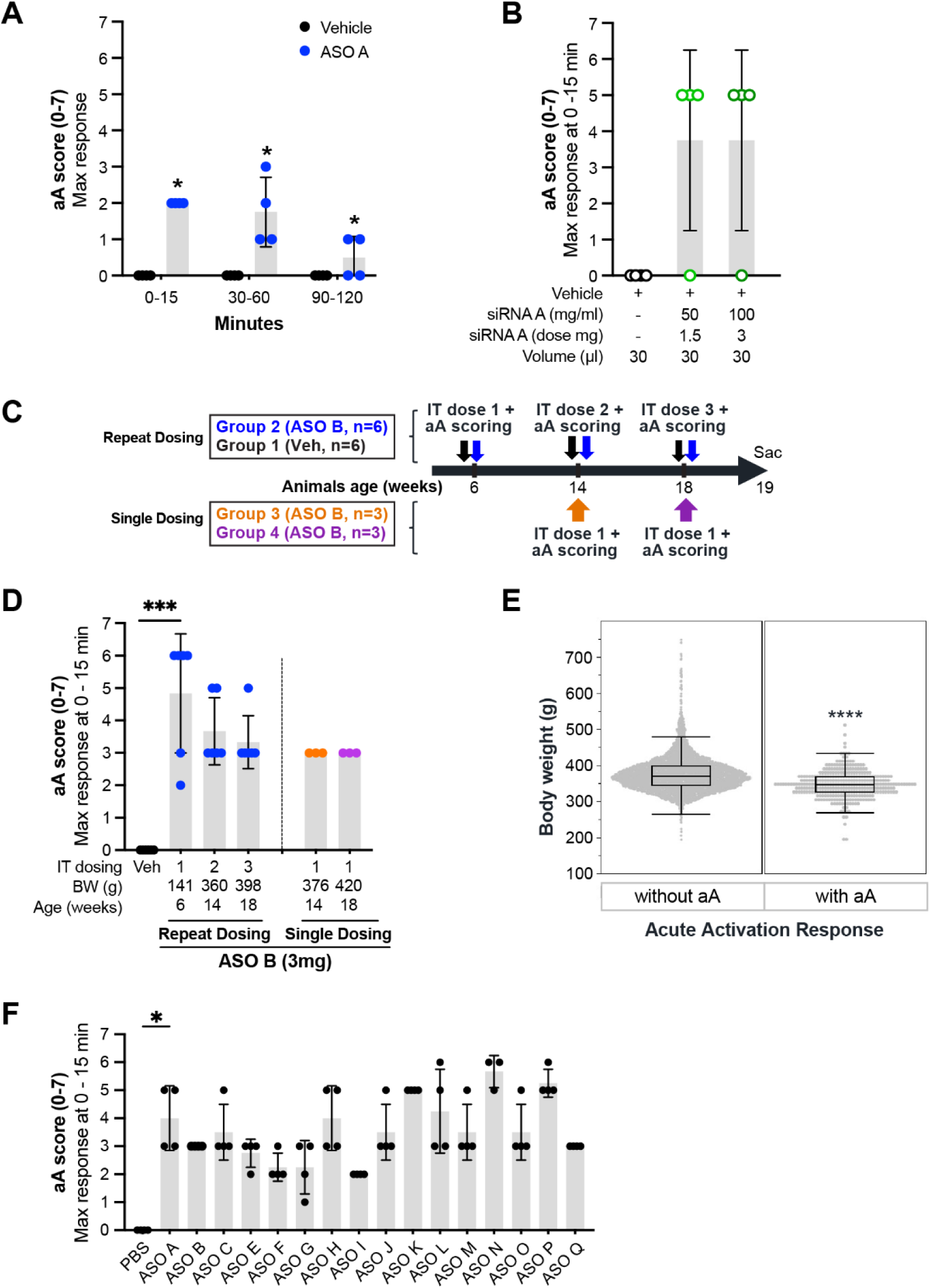
Body-weight dependent, transient acute neuronal activation (aA) response after intrathecal oligonucleotide injection. **(A)** Time-course of acute neuronal activation (aA) response in male rats (body weight = 261 ± 5 g) after single intrathecal (IT) administration of 3 mg ASO A in 30 µl (100 mg/ml, 14.1 mM) (blue circles, n= 4) or 30 µl of artificial cerebrospinal fluid (Vehicle, black circles, n= 4). The aA scores for each group (ASO A or Vehicle), were divided into three-time blocks (0-15 minutes, 30-60 minutes, and 90-120 minutes), revealing a significant increase in response during the first hour, which then decreases over the subsequent hour. Data are expressed as mean ± s.d. The statistical significance was determined by the Mann-Whitney test *P ≤ 0.05; ** P ≤ 0.01 *** P ≤ 0.001; **** P ≤ 0.0001 *vs* vehicle (aCSF) at each time point **(B)** Acute neuronal activation maximum response in male rats (body weight = 384 ± 32 g) after 0-15 min of a single IT administration of 1.5 g (light green circles, n= 4) or 3 mg (dark green circles, n=4) of siRNA A or PBS (Vehicle, black open circles, n= 4). Data are expressed as mean ± s.d. The statistical significance was determined by the Kruskal-Wallis test. *P ≤ 0.05; ** P ≤ 0.01 *** P ≤ 0.001; **** P ≤ 0.0001 *vs* vehicle (PBS). **(C)** Illustration of the dosing experimental design for evaluating aA in both single and repeat dosing paradigms. Black and blue arrows denote animals receiving repeated dosing with PBS (Vehicle) or ASO B (100 mg/ml, 14.1 mM), respectively. Orange and purple arrows represent animals receiving a single dose of ASO B at specified times. **(D)** Acute neuronal activation maximum responses in male rats measured within the first 15 minutes after repeat (Blue circles, n= 6) or single (orange and purple circles, n= 4) IT administration of ASO B (3 mg in 30 µl) show that aA scores remain consistent across three doses (blue) compared to a single dose (orange and purple). The vehicle repeat dose group (black circles, n=6 males) showed no significant phenotype. Data are expressed as mean ± s.d. The statistical significance was determined by the Kruskal-Wallis test. *P ≤ 0.05; ** P ≤ 0.01 *** P ≤ 0.001; **** P ≤ 0.0001 *vs* ASO B, IT dosing 1 **(E)** Occurrence of acute neuronal activation response after ASO administration in male rats and the body weight. The figure shows that the occurrence of aA is more likely in treated animals with lower body weights. Animals with body weight greater than 450 g are very unlikely to experience acute neuronal activation (score ≥ 5). Data are expressed as mean ± s.d. The statistical significance was determined by parametric unpaired t-test *P ≤ 0.05; ** P ≤ 0.01 *** P ≤ 0.001; **** P ≤ 0.0001. **(F)** Acute neuronal activation maximum response induced in male rats (n=4, body weight 294 ± 13 g) by sixteen ASOs same chemistry (5-10-5 MOE gapmer, mix backbone [PO/PS], Supplementary table 1) and different sequences. Data are expressed as mean ± s.d. The statistical significance was determined by the Kruskal-Wallis test. *P ≤ 0.05; ** P ≤ 0.01 *** P ≤ 0.001; **** P ≤ 0.0001 *vs* ASO A. Abbreviations: IT=intrathecal; Sac=sacrificed; aA= acute neuronal activation ; aCSF: artificial cerebrospinal fluid.

Double-stranded siRNAs have also been reported to exhibit similar acute neuronal activation responses (11, 23). To quantify this, we used the same scoring system and treated rats via IT administration with either 1.5 or 3 mg of siRNA A. This siRNA was previously optimized for IT delivery in rats, but it was not evaluated for acute tolerability (32). The acute neuronal activation response of the siRNA was severe at both the 1.5 and 3 mg dose levels (**Figure 1B**), confirming this behavioral response can occur with both single-stranded ONs and double-stranded ONs.

To test another ASO sequence and to explore if the acute neuronal activation response changed with repeat dosing, we administered ASO B (**Supplementary Table 1**) to rats in a repeat dosing paradigm at ages 6, 14, and 18 weeks and evaluated the acute neuronal activation response after each dose (**Figure 1C**). The peak response was after the first administration. The animals that received the repeat dose regimen exhibited a non-significant decrease in response after the second and third doses (**Figure 1D**). As the rats gain weight from 6-18 weeks of age, to control for body weight changes, two independent, body-weight-matched groups were injected with a single dose at 14 and 18 weeks, respectively. The animals in the single-dosed groups (**Figure 1D, orange and purple circles**) displayed the same acute neuronal activation response as the corresponding age groups in the repeat-dosing paradigm (**Figure 1D, blue circles**). These results suggest that acute neuronal activation responses likely do not change with repeat ASO administration, but that the size of the animal may contribute to the magnitude of the response.

To determine if body weight was a factor, we reviewed more than two thousand rat entries from screening studies, with ∼640 different ASOs, yielding an 18.5% incidence (455 occurrences out of 2464 entries) of severe acute neuronal activation responses (vocalization/seizures) after intrathecal administration of ASO into male rats of varying size. The acute neuronal activation response was more likely to occur in animals with lower body weights (**Figure 1E**) and did not occur in very large animals (>450 grams). No clear patterns for sequence dependence emerged from the evaluation of the database (data not shown). Taken together, the acute neuronal activation response is variable at this dose level (3 mg), and body weight is an important covariate of the acute neuronal activation response that should be controlled for.

To directly test whether the acute neuronal activation effect is sequence-dependent, fifteen ASOs with identical chemistry as ASO A but unique sequences were tested (**Supplementary Table 1**). The acute neuronal activation response ranged from 2 to 6, with no clear sequence dependence (**Figure 1F**).

There are two behaviorally distinct phenotypic responses in rats with phosphorothioate ASOs following IT administration. One is the acute neuronal activation response described here, and the other is the acute sedation response. The acute sedation response is highly sequence-dependent, and peaks 3 hours after dosing, resolving by 24 hours (13). We evaluated our database of over 600 ASOs that induced an acute sedation phenotype, characterized by an ‘inhibitory’ response as measured by the acute sedation score (**Supplementary Table 2,** (13)), and there was no enrichment of ASOs that exhibit an acute neuronal activation response (**Supplementary Figure 1**), suggesting no connection between the occurrence of acute sedation response and the presence of acute neuronal activation response. Consistent with the temporal disconnect between the peak and resolution of the activation and sedation responses, these two acute responses are likely driven by distinct mechanisms.

### Acute neuronal activation response occurs following the administration of oligonucleotides into the CSF of mice and non-human primates

Many preclinical studies evaluating ON for the treatment of neurological diseases utilize mouse models. IT delivery is difficult in mice, thus intracerebroventricular (ICV) delivery to the CSF is often used (33–35). In the present study, we designed an ICV-specific score, which focuses exclusively on activation behaviors and can be used to evaluate the acute neuronal activation behaviors based on the severity (**Table 2**).

To determine the translatability across delivery routes and species, we evaluated the acute neuronal activation response in mice (**Figure 2A-C**). First, a dose-response study using ASO A was conducted in male and female mice. We evaluated the behavior of the mice for two hours after dosing. The acute neuronal activation response was observed starting at 700 µg. The maximum responses were observed within the first 15 to 30 min after the administration, but after 1 hour most animals behaved similarly to their control littermates (**Figure 2A).** Female mice exhibited more pronounced signs of acute neuronal activation compared to males, consistent with their smaller body size (female body weight: 18 ± 1 g; male body weight: 23 ± 2 g) (**Figure 2B, left and middle panels**). A similar sex- and size-related difference was observed in rats. Age-matched female rats, which were smaller (average body weight: 204 ± 4 g), displayed a stronger acute neuronal activation response (**Supplementary Figure 2**) than their larger male counterparts (average male body weight: 261 ± 5 g **in Figure 1A**) under the same dosing paradigm.

**Figure 2.**
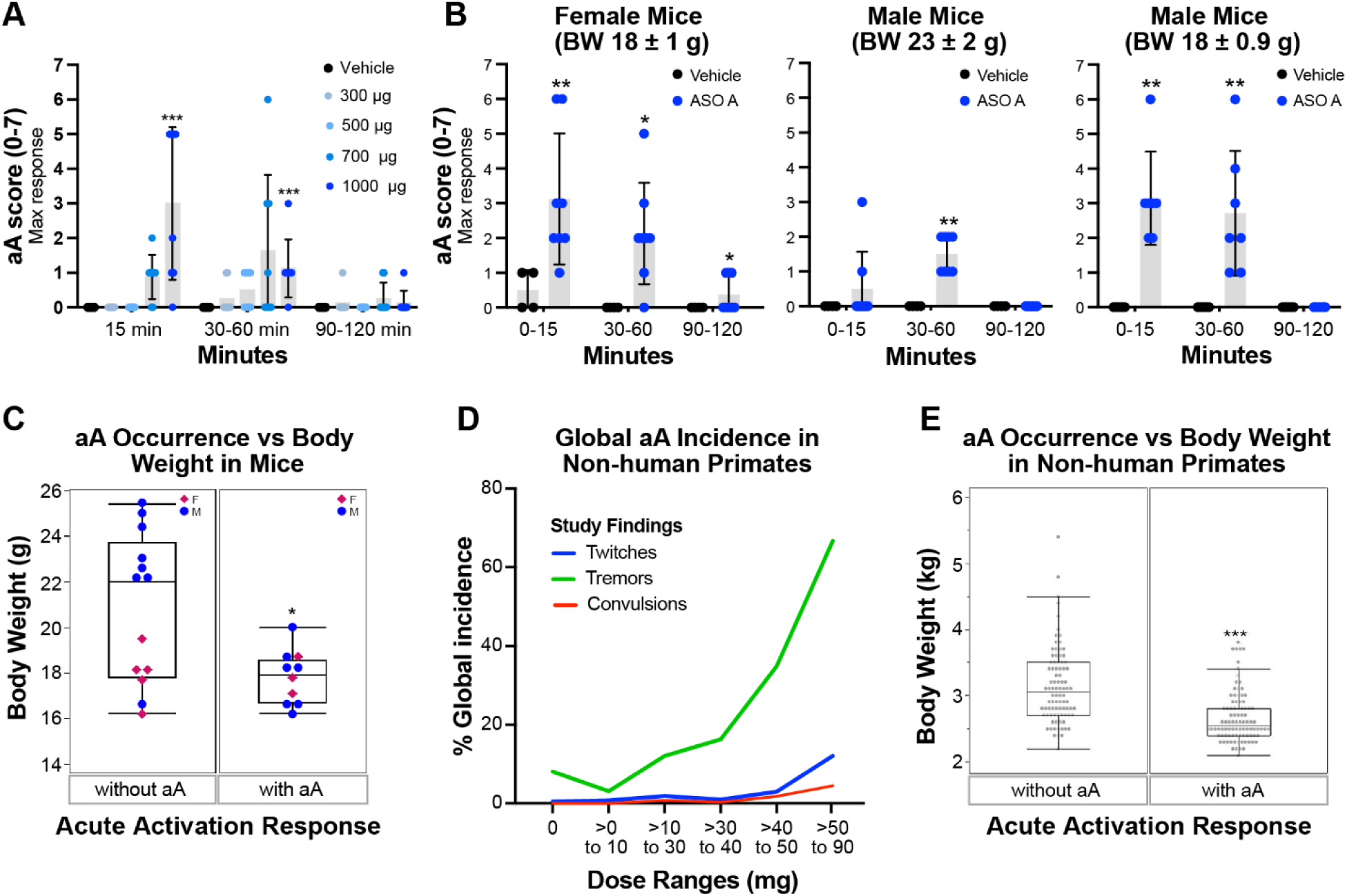
Acute neuronal activation response occurs following the injection of oligonucleotides into the CSF of mice and non-human primates. (A-B) Time-course of acute neuronal activation maximum response is divided into three-time blocks (0-15 minutes, 30-60 minutes, and 90-120 minutes) after 10 µl of a single intracerebroventricular (ICV) administration in mice. **(A)** Dose-response of acute neuronal activation maximum response after 300, 500, 700 and 1000 µg ASO A (n= 4 females, BW= 17.7 ± 1 g and n≥ 4 males, BW= 21.6 ± 1.5 g) or vehicle (aCSF, black circles, n=4). **(B)** Acute neuronal activation maximum response after 1000ug of ASO A (100mg/ml, 14.1 mM) in females (n=8) with an average body weight of 18 ± 1 g (left panel) and age-matched males (n=8) with a body weight of 23 ± 2 g (middle panel). Acute neuronal activation maximum response in males (n=7) with similar body weight as females, 18 ± 0.9 g (right panel). These data reveal a significant body weight-dependent increase in response at 0-15 min that can last up to an hour, which then decreases over the subsequent hour compared with the vehicle group. Each symbol represents an individual animal. The statistical significance was determined by the Mann-Whitney test *P ≤ 0.05; ** P ≤ 0.01 *** P ≤ 0.001; **** P ≤ 0.0001 *vs* vehicle (aCSF) at each time point. **(C)** Occurrence of acute neuronal activation response and mice body weight; each dot represents a unique animal. Mice were more likely to experience at least one acute neuronal activation event (score ≥3) when body weights were lower than 20 g. The statistical significance was determined by parametric unpaired t-test *P ≤ 0.05; ** P ≤ 0.01 *** P ≤ 0.001; **** P ≤ 0.0001. **(D)** The dose ranges used in NHP, when plotted against the percent global incidence of acute neuronal activation response, illustrate how the severity of responses—ranging from less severe (twitches) to moderate (tremors) and most severe (convulsions)—increased with higher doses. 0 mg represents vehicle control (aCSF). **(E)** Occurrence of acute neuronal activation response and adult nonhuman primate body weight; each dot represents a unique animal. Nonhuman primates were more likely to experience at least one aA event with body weights lower than 4 kg. The data is expressed as the mean ± s.d. The statistical significance was determined by parametric unpaired t-test *P ≤ 0.05; ** P ≤ 0.01 *** P ≤ 0.001; **** P ≤ 0.0001. Abbreviations: ICV=intracerebroventricular; aA= acute neuronal activation; aCSF=artificial cerebrospinal fluid.

To directly assess the contribution of body weight to acute neuronal activation responses in mice, we performed the same experiment using smaller males with body weights between 17 and 18 g. As expected, these smaller males exhibited more severe acute neuronal activation responses, comparable to those of females in the same body weight range (**Figure 2B, right panel**). Similar to rats, the occurrence of the acute neuronal activation phenotype in mice was influenced by body weight, as animals weighing over 20 g were less likely to exhibit a severe acute neuronal activation response compared to smaller animals (**Figure 2C**). Additionally, there was a strong inverse correlation between the severity of acute neuronal activation and body weight (p = 0.0011, **Supplementary Figure 3A**) and in all cases, the acute neuronal activation response was resolved by 120 minutes regardless of body weight (**Supplementary Figure 3B-D**). Overall, these results suggest that mice also display transient acute neuronal activation responses, and that body weight is an important covariate in the severity of the ASO-induced acute neuronal activation response across species.

Acute neuronal activation response like that in rats has also been observed in preclinical studies in non-human primates (NHP) after intrathecal administration of ASOs formulated in aCSF. A retrospective analysis of the incidence of acute neuronal activation responses from our internal database indicates that the most common observation is tremors, and the incidence of convulsions is rare (**Figure 2D**). Furthermore, the query suggested that the incidence and severity of acute neuronal activation responses in NHP increases in a dose-dependent fashion, with acute neuronal activation being observed frequently at doses above 50 mg (human equivalent dose of 500 mg). Similar to rats, the occurrence of the acute neuronal activation phenotype in NHP was influenced by body weight, as animals over 4 kg were less likely to present with an acute neuronal activation response than smaller animals (**Figure 2E**). The onset of the acute neuronal activation phenotype in NHPs is typically immediately after dosing or during recovery from anesthesia with full resolution after approximately 120 min with no apparent sequelae. The signs and behaviors observed include tremors, muscle twitches, clonic or tonic movements of the limbs, salivation, self-biting, opisthotonos, and/or generalized convulsions or seizures. Diazepam administered to NHP has helped to alleviate these signs and behaviors. If severe enough, this acute neuronal activation response can be a dose-limiting finding in NHP.

To date, a scoring system that accurately captures this activation phenotype in NHP has not been described. Consequently, we developed an acute neuronal activation scoring system that classifies the signs and behaviors based on their severity. A score is assigned to each set of behaviors, ranging from no relevant clinical findings (0) to requiring euthanasia (7) as described in **Table 3**.

### Increasing the divalent cations-to-ASO ratio effectively mitigates the acute neuronal activation response

ON-induced seizures have been reported to be mitigated by supraphysiological levels of calcium or magnesium (23). We hypothesize that the acute neuronal activation response observed here is driven by ON-mediated chelation of endogenous divalent cations in the intrathecal space, particularly when ASO concentrations are elevated in the CSF immediately after dosing. Supporting this, oxalic acid, which binds divalent cations like calcium, magnesium, and zinc, reduces their bioavailability by forming insoluble oxalate precipitates (36, 37). It causes muscle tremors, ataxia, tetany, salivation, and convulsions in sheep and cattle (2–6 h post-oral administration) (36) and has been shown to induce seizures immediately following 15–20 mM intrathecal administration in mice and rats (our observation in an independent study, data not shown). In all these cases the hypo-divalent cationic state likely triggers local CNS cell activation, consistent with symptoms of severe hypomagnesemia and hypocalcemia, including tremors, cramps, twitching, and seizures (18, 38, 39).

Due to the low fixed volumes required for IT dosing, our typical formulations in preclinical safety studies have high ON concentrations relative to the physiological concentrations of divalent cations used in the dosing solution. To test our hypothesis, we evaluated the acute neuronal activation response in rats with different concentrations of divalent cations, varying the molar ratio of divalent cations-to-ON in the dosing solution. Since we hypothesized that ON deplete endogenous cations in the CSF and parenchyma when there are insufficient cations in the dosing solution to saturate ON-cation binding, we focused on investigating acute neuronal activation responses were affected by the molar divalent cations-to-ON ratio rather than by total dose or total cation levels. This approach will also allow for translation across species, as different dosing volumes are used for different species. Here, ASO A was formulated with divalent cations-to-ASO ratios from 0.16:1 (levels in aCSF; Calcium = 1.4 mM and Magnesium = 0.8 mM) to 0.3:1, 0.7:1, 1.3:1, 1.7:1, 2.3:1 and 3:1. The divalent cations-to-ASO ratio was adjusted by adding magnesium, calcium or the combination of both proportional to their physiological ratio (1.75 Calcium: 1 Magnesium, **Supplementary Tables 3-5**). Intrathecal administration with ASO A showed a typical acute neuronal activation response pattern compared with the control group (aCSF) (**Figure 3**). In rats treated with the same dose, but the divalent cations-to-ASO ratio was adjusted by adding magnesium to the dosing solution, the acute neuronal activation responses were substantially reduced at all time points when the divalent cations-to-ASO ratios were ≥ 2.3:1 (**Figure 3A**). Interestingly, when the divalent cations-to-ASO ratio was adjusted by adding calcium to the ASO formulation, there was a robust mitigation effect of the acute neuronal activation response at 15 minutes post-dosing starting at a divalent cations-to-ASO ratio of 0.7:1, reducing the score to control group levels (aCSF) (**Figure 3B, left panel**). However, at approximately 30 minutes, animals in all groups receiving ASO dosing solution supplemented with calcium (but not calcium alone) began exhibiting pronounced activation responses. These included shaking, involuntary whole-body movements (such as hopping in place or moving around the cage), and mild vocalizations (low-pitched squeaking). Additionally, they displayed hyperreactivity to stimuli (e.g., cage mate movements, sounds), as well as defensive or aggressive behaviors. These responses persisted for up to 120 minutes.

**Figure 3.**
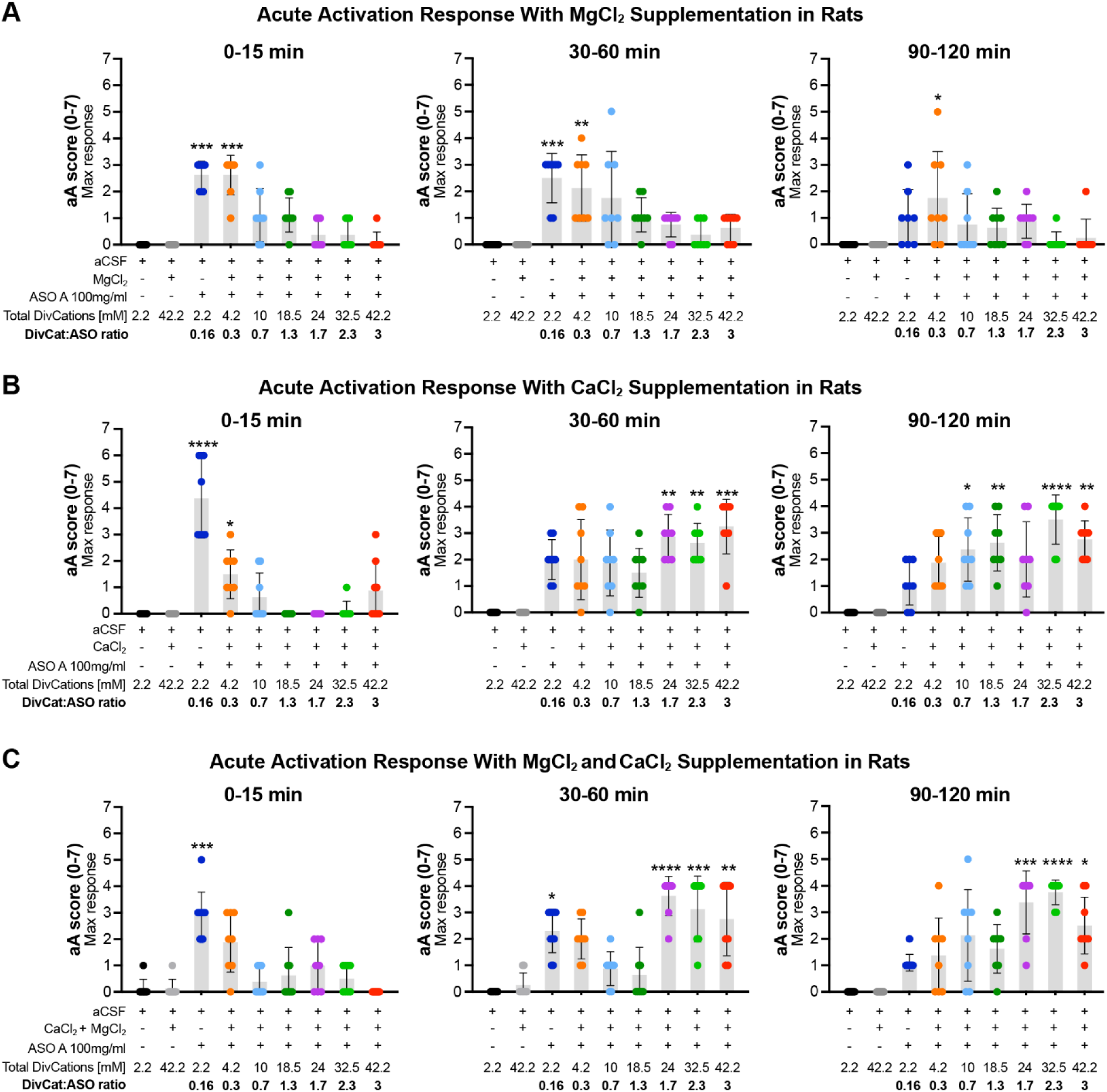
Increasing the divalent cation-to-ASO ratio mitigates acute neuronal activation response. **(A)** Acute neuronal activation (aA) maximum response in rats plotted in three-time blocks; 0-15 minutes (left); 30-60 minutes (middle); and 90-120 minutes (right) after intrathecal 3 mg dose administration of ASO A (100 mg/ml, 14.1 mM) formulated in aCSF with or without increasing millimolar concentrations of magnesium supplementation, resulting in six different DivCat: ASO ratio from 0.3:1 to 3:1 (n≥ 4 females, BW= 225 ± 8 g and n≥ 4 males, BW= 286 ± 11 g). The data shows a significant reduction of aA response in a ratio-dependent manner over the 2 hours. **(B)** Acute neuronal activation maximum response in rats plotted in three-time blocks; 0-15 minutes (left); 30-60 minutes (middle); and 90-120 minutes (right) after intrathecal 3 mg dose administration of ASO A (100 mg/ml, 14.1 mM) formulated in aCSF with or without increasing millimolar concentrations of calcium supplementation, resulting in six different DivCat: ASO ratio from 0.3:1 to 3:1 (n≥ 4 females, BW= 220 ± 10 g and n≥ 4 males, BW= 285 ± 10 g). The data shows a significant reduction of aA response in a ratio-dependent at only 0-15 min time point with a clear, sustained, and significant rebound acute activation effect at all tested ratios from the 30 min up to the 2 hours. **(C)** Acute neuronal activation maximum response in rats plotted in three-time blocks; 0-15 minutes (left); 30-60 minutes (middle); and 90-120 minutes (right) after intrathecal 3 mg dose administration of ASO A (100 mg/ml,14.1 mM) formulated in aCSF with or without increasing millimolar concentrations of magnesium plus calcium supplementation (kept at their physiological ratio), resulting in six different DivCat: ASO ratio from 0.3:1 to 3:1 (n≥ 4 females, BW= 224 ± 8 g and n≥ 4 males, BW= 294 ± 8 g). The data shows a significant reduction of aA response in a ratio-dependent at only 0-15 min time point with a clear, sustained, and significant rebound acute activation effect at the 3 highest ratios from the 30 min up to the 2-hours. Data are expressed as mean ± s.d. Each symbol represents an individual animal. The significant difference was determined by the Kruskal-Wallis test. *P ≤ 0.05; ** P ≤ 0.01 *** P ≤ 0.001; **** P ≤ 0.0001 *vs* vehicle (aCSF). Abbreviations: Aa, acute neuronal activation; aCSF, artificial cerebrospinal fluid; DivCat, Divalent cations.

Thus, the mitigation effect of calcium supplementation was only transient and ultimately led to an extended activation response (**Figure 3B, middle and right panels**). Similar effects were observed in rats treated with ASO formulations where divalent cation-to-ASO ratios were adjusted using a combination of magnesium and calcium (proportional to their physiological levels) (**Figure 3C**). The complete time course of the acute neuronal activation responses and the calculated area under the curve confirm that increasing the divalent cations-to-ASO ratio by adding magnesium to the formulation suppresses the acute neuronal activation phenotype in rats in a ratio-dependent manner (**Supplementary Figure 4**), whereas formulations supplemented with calcium only transiently mitigated and then prolonged the acute neuronal activation response.

To test whether the observed rebound effect with calcium supplementation was related to the hypertonic nature of this formulation (∼620 mOsm/L), we intrathecally delivered ASO A formulated in aCSF in which the tonicity was controlled by reducing the sodium concentration achieving isotonicity (260-340 mOsm /Kg) commensurate with ASO and the increase in calcium concentration (dACSF) (**Supplementary Table 6**). However, the same rebound effect was observed 30 min after the administration, ruling out that the rebound in response was related to the tonicity of the dosing solution (**Supplementary Figure 5**).

No other acute safety endpoints were altered with this supplementation and all animals gained body weight like control animals (2 weeks after dosing) (**Supplementary Figure 6**).

To determine if this mitigation approach is broadly applicable, we formulated another ASO, ASO C (**Supplementary Table 1**), with or without cation supplementation by adding magnesium to the ASO formulation. As expected, the acute neuronal activation responses were significantly mitigated with a total divalent cations-to-ASO ratio of 2.3:1 (**Supplementary Figure 7**). The acute neuronal activation response for ASO A could also be mitigated with magnesium supplementation under both hypertonic and isotonic conditions (**Supplemental Figure 8, Table Supplementary 7**).

### No identified consequences of modifying the divalent cations-to-ASO ratio with magnesium supplementation

Next, we investigated whether modifying the molar divalent cation-to-ASO ratio with magnesium supplementation by a maximum of 3:1 is tolerated. Animals from our previous experiment were followed for 8 weeks and then sacrificed to measure neuroinflammatory markers by qPCR and immunohistochemistry. The body weight was comparable between all groups (**Figure 4A-C**) and no physical signs of illness were observed. ASO A induced a slight increase in mRNA levels of *Gfap* and *Aif1* in the cortex and spinal cord eight weeks after dosing compared with the vehicle groups. Magnesium supplementation did not change the neuroinflammatory markers quantified by qPCR (**Figure 4D-G**) or immunohistochemistry (**Figure 4H-K**).

**Figure 4.**
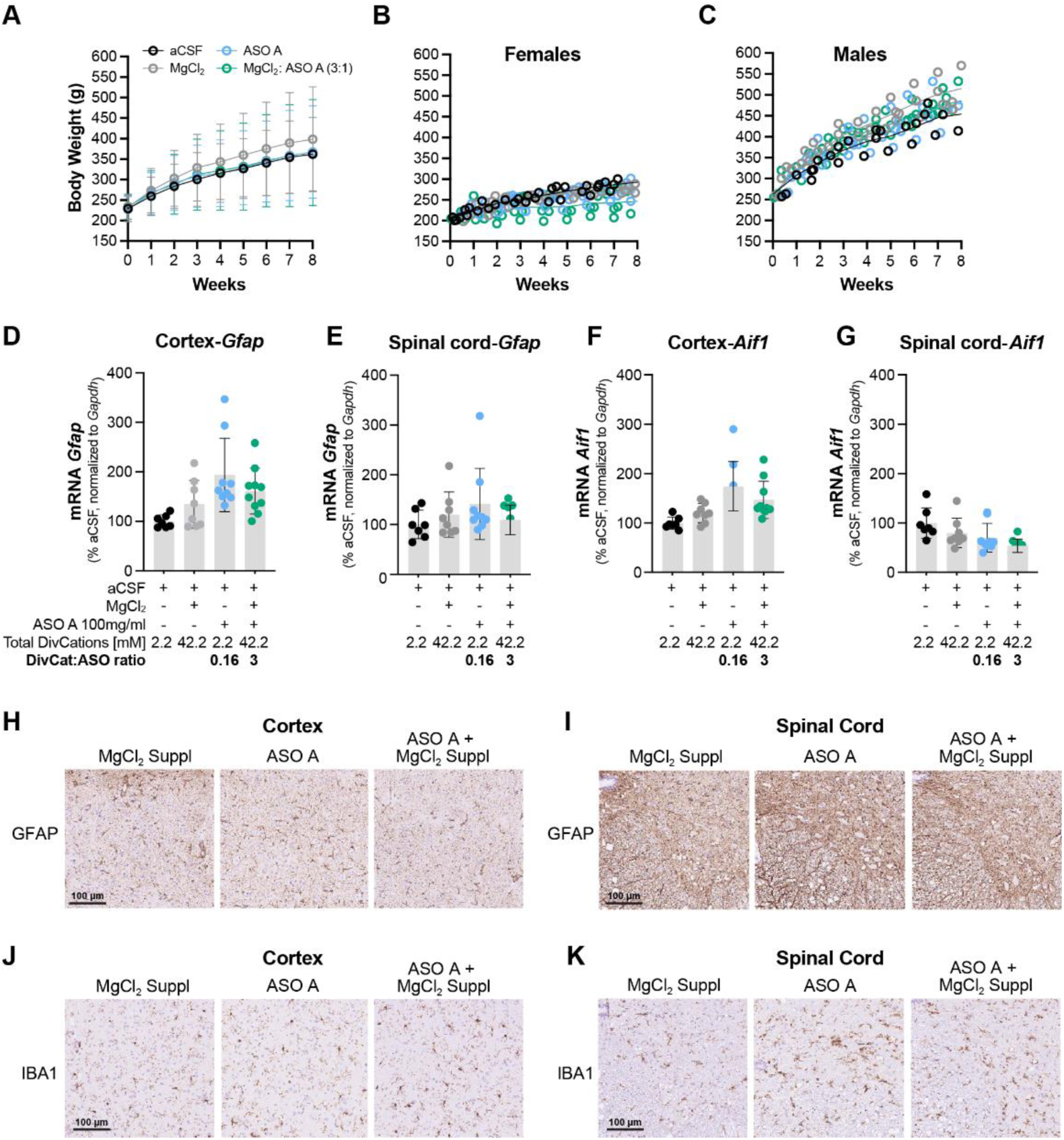
No identified consequences of modifying the divalent cation to ASO ratio with magnesium supplementation. **(A-C)** Body weight gain over 8 weeks (**A**) after intrathecal 3 mg dose administration of ASO A (100 mg/ml, 14.1 mM) with or without magnesium supplementation (40 mM) to a DivCat: ASO ratio of 3:1 formulated in artificial cerebrospinal fluid (aCSF). Note normal body weight gain is achieved in females (**B**) and males (**C**). (**D-G**) mRNA expression of *Gfap* (**D, E**) or *Aif1* (**F, G**) in the cortex (**D, F**) or spinal cord (**E, G**) after 8 weeks of intrathecal 3 mg dose administration of ASO A (100 mg/ml,14.1 mM) with or without magnesium supplementation (40 mM) to a DivCat: ASO ratio of 3:1 formulated in aCSF. (**H-K)** Representative IHC images of cortex or spinal cord with GFAP (**H, I**) or Iba1 (**J, K**) staining respectively, after 8 weeks of intrathecal 3 mg dose administration of ASO A (100 mg/ml, 14.1 mM) with or without magnesium supplementation (40 mM) to a DivCat: ASO ratio of 3:1 formulated in aCSF. Each symbol represents an individual animal. Data are expressed as mean ± s.d. (n≥ 4 females, BW= 204 ± 4 g, and n≥ 4 males, BW= 261 ± 5 g). The significant difference was determined as follows; RM two-way ANOVA (A-C) or One-way ANOVA (D-G) followed by Tukey’s post hoc test. *P ≤ 0.05; ** P ≤ 0.01 *** P ≤ 0.001; **** P ≤ 0.0001 comparing vehicle (aCSF) *vs* ASO or comparing ASO with or without magnesium supplementation. No significant differences were found. Abbreviations: Aa, neuronal activation response; aCSF, artificial cerebrospinal fluid; DivCat, Divalent cations; MgCl_2_ suppl, Magnesium supplementation. Scale= 100 µm.

Overall, these experiments suggest that formulations of ASO with up to 40 mM of magnesium supplementation in aCSF, at a divalent cation-to-ASO ratio of 3:1, safely mitigate the neuronal activation response, without sequelae or change to other tolerability endpoints.

### Magnesium supplementation does not alter oligonucleotide activity, tissue uptake, and distribution

To assess the impact of magnesium supplementation on the efficacy and potency of ASO A, a human *ATXN3* targeting ASO, we treat transgenic mice expressing the full-length human *ATXN3* gene with ICV administration of the calculated ED80 with or without supplementation with 40 mM magnesium (**Supplementary Table 8**). Tissues were harvested 2 weeks after dosing for analysis. ASO formulations with or without magnesium supplementation similarly suppressed target mRNA levels in the cortex (**Figure 5A**) and spinal cord (**Figure 5B**). Similarly, the ASO concentration in tissues and ASO localization in tissue (measured by IHC ASO staining) were comparable between conditions in the cortex (**Figure 5C and E**) and spinal cord (**Figure 5D and F**).

**Figure 5.**
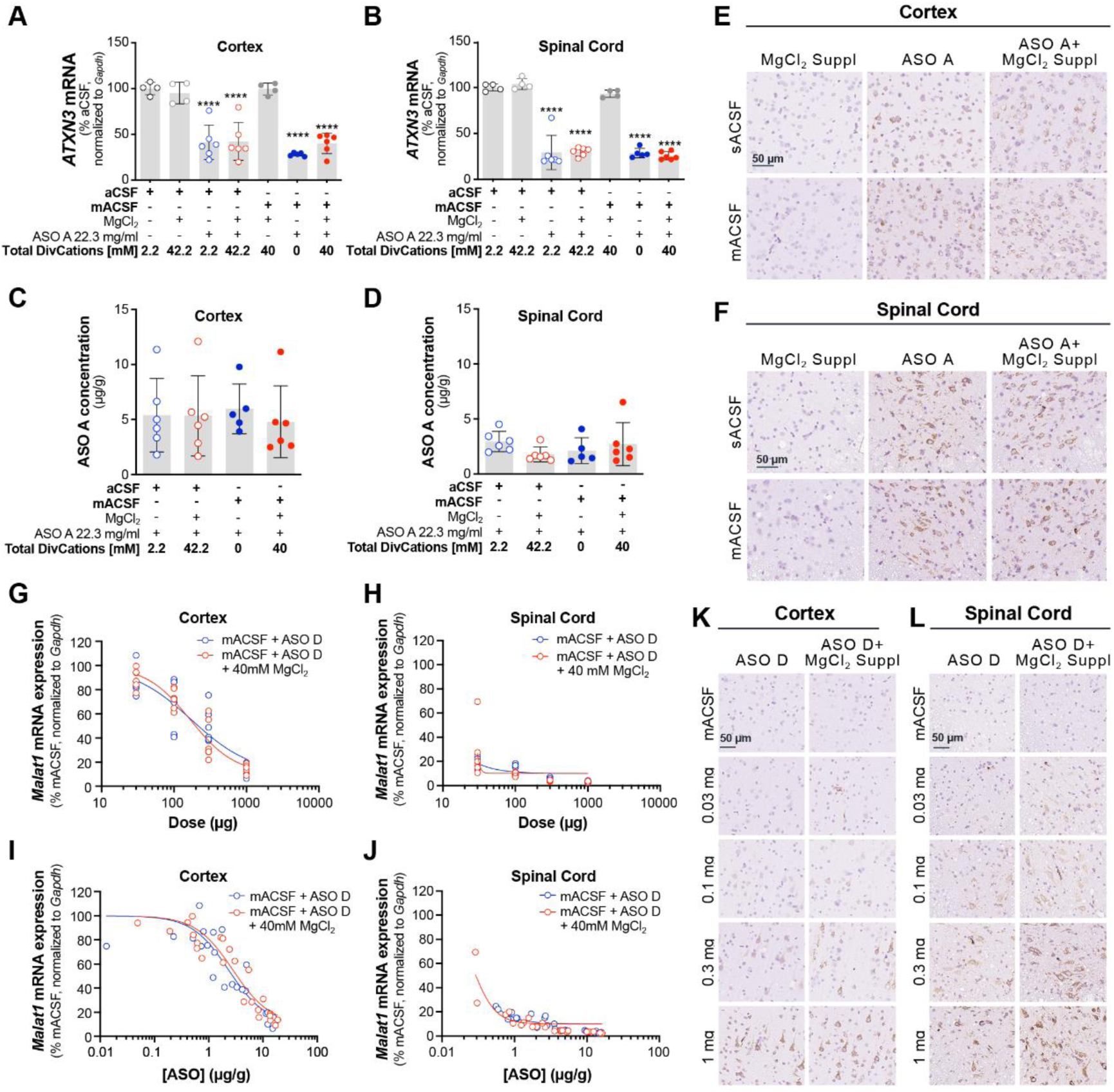
Magnesium supplementation does not alter oligonucleotide activity, tissue uptake, and distribution. **(A-F)** *ATXN3* mRNA expression levels in cortex (**A**) and spinal cord (**B**) and ASO tissue concentration in the cortex (**C**) and spinal cord (**D**) in transgenic mice (n≥2 female, BW=18 ± 1 g; n≥2 male mice, BW= 21± 1 g) after 2 weeks of intracerebroventricular (ICV) injection of the calculated ED80 (223 µg) administration of ASO A (22.3 mg/ml, 3.14 mM) with or without magnesium supplementation (40 mM) formulated in 10 µl of standard artificial cerebrospinal fluid (aCSF) or a modified aCSF (mACSF; magnesium and calcium removed). Representative IHC images of ASO staining in the cortex (**E**) or spinal cord (**F**) are shown. **(G-L)** The dose-response curve of mRNA expression of rat *Malat1* in the cortex (**G**) and spinal cord (**H**) (n=6 males, BW= 305 ± 9 g). Rat *Malat1* mRNA expression plotted against ASO concentration in cortex (**I**) and spinal cord (**J**). Representative IHC images of ASO staining in the cortex (**K**) or spinal cord (**L**) after 2 weeks of intrathecal dose-response administration of ASO D (0.03, 0.1, 0.3, 1 mg in 30 µl) with or without magnesium supplementation (40 mM) formulated in a modified aCSF (mACSF; magnesium and calcium removed). Note how in rats and mice, ASO activities remain unaffected in any condition when magnesium is supplemented. Each point represents an individual animal. Data is expressed as follows: Mice data is expressed as the mean ± s.d. of the percent of aCSF *ATXN3* mRNA levels normalized to *Gapdh* (A and B) or the ASO concentration (µg/µl) (C and D) against the different groups. The rat dose-response data are expressed as a percent of aCSF *Malat1* mRNA levels normalized to *Gapdh* against log10 of the dose (µg) (G and H); the concentration-response curve is plotted as a percent of aCSF *Malat1* mRNA levels normalized to *Gapdh* against log10 of the tissue concentration (µg/µl) (I and J). ED_50_ and EC_50_ were calculated using the Motulsky: Agonist vs. response -- Variable slope (four parameters) and the F-test method was used to compare the difference between groups. The significance difference was determined by one-way ANOVA followed by Tukey’s post hoc test. *P ≤ 0.05; ** P ≤ 0.01 *** P ≤ 0.001; **** P ≤ 0.0001 *vs* the corresponding vehicle. No differences were found when comparing vehicle or ASO with and without magnesium supplementation. Abbreviations; ICV, intracerebroventricular; aCSF, artificial cerebrospinal fluid; mACSF, modified artificial cerebrospinal fluid; MgCl_2_ suppl, Magnesium supplementation. Scale= 50 µm.

To further characterize the effect of magnesium supplementation on ASO activity and uptake, we used ASO D, an antisense oligonucleotide targeting rat *Malat1* RNA. This ASO has been previously characterized for its dose-responsive *Malat1* RNA reduction and accumulation in rat CNS tissue (6). Thus, rats received a 30 µl intrathecal administration of 0.03 mg, 0.1 mg, 0.3 mg, or 1 mg with or without 40 mM of magnesium supplementation (**Supplementary Table 9**), and tissues were harvested 2 weeks after dosing for analysis. The ASO formulations with or without magnesium demonstrated similar decreases in *Malat1* mRNA levels in a dose-dependent manner with comparable ED_50_ values between conditions; ASO D in mACSF: cortex = 174.9 µg and ASO D + 40 mM magnesium in mACSF: cortex = 168.4 µg (F 1.325, DFn 2, DFd 39; P=0.2775 vs ASO in mACSF) (**Figure 5G and H**). Consistently, the tissue ASO concentration-response curve overlapped between groups generating a comparable EC_50_ between conditions; ASO D in mACSF: cortex = 2.3 µg (95% CI: 1.699 to 3.216 µg) and ASO D + 40 mM magnesium in mACSF: cortex = 2.792 µg (95% CI: 2.104 to 3.628 µg) (**Figure 5I and J**). In line with these findings, ASO concentration increased in a dose-dependent manner formulated with or without magnesium supplementation (**Supplementary Figure 9**). Moreover, staining for the presence of ASO using immunohistochemistry (IHC) demonstrates indistinguishable ASO uptake and localization in the cortical (**Figure 5K**) and spinal cord neurons with or without magnesium supplementation (**Figure 5L**). These data confirm that supplementation of the ASO formulations with up to 40 mM of magnesium does not impact the efficacy, distribution, or uptake of ASO in the CNS of rodents.

### The divalent cation-to-ASO ratio is the primary factor responsible for mitigating the acute neuronal activation response

Finally, to verify that adjusting the ratio of divalent cations-to-ASO effectively mitigates the neuronal activation response, we increased the ASO dose volume from 30 µl to 100 µl while maintaining a total dose of 3 mg (**Supplementary Table 10**). Increasing the dose volume decreases the concentration of ASO from 14.1 mM (100 mg/ml) to 4.22 mM (30 mg/ml) while the cation concentration in the aCSF is constant at 2.2 mM total divalent cations. This adjustment (increasing the dose volume) increases the ratio of total divalent cations (in aCSF) to ASO from 0.16:1 to 0.52:1, and this alone diminishes the acute neuronal activation score (**Figure 6A-C**). Further attenuation of the response can be achieved by adding as little as 10 mM magnesium to the 100 µl dosing solution to create a 3:1 divalent cation-to-ASO ratio, and this has been replicated with a second ASO (**Figure 6B, C**). These data confirm that at a given dose level of ASO, the ratio of cations to ASO in the dosing solution drives the response, rather than the total amount of cations or ASO administered.

**Figure 6.**
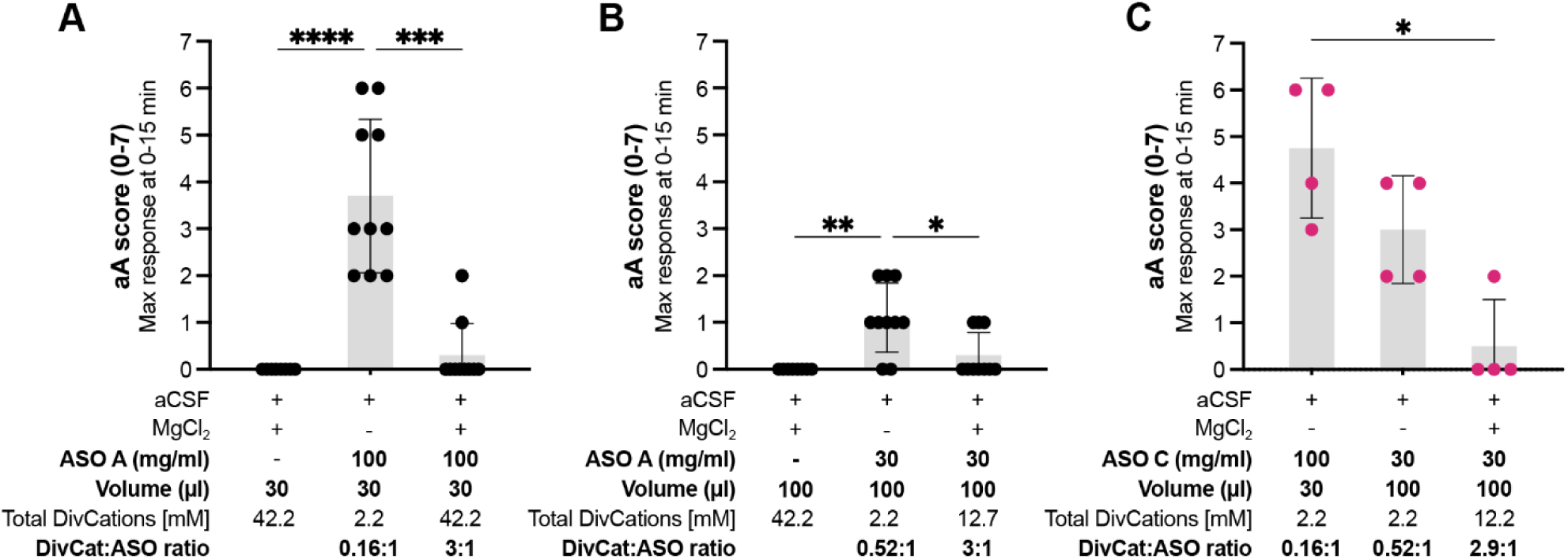
The divalent cation to ASO ratio is primarily responsible for the mitigation of the acute neuronal activation response. **(A)** Acute neuronal activation (aA) maximum response measured at 0-15 min after 30 µl intrathecal (IT) administration of aCSF with magnesium supplementation (40 mM), or 3 mg dose of ASO A (100 mg/ml; 14.1 mM) alone or 3 mg dose of ASO A (100 mg/ml; 14.1 mM) with magnesium supplementation (40 mM) to a DivCat: ASO ratio of 3:1. Note that magnesium supplementation shows significant aA mitigation. **(B)** Acute neuronal activation maximum response after 100 µl intrathecal administration of aCSF with magnesium supplementation (40 mM), or 3 mg dose of ASO A (30 mg/ml; 4.22 mM) alone or 3 mg dose of ASO A (30 mg/ml; 4.22 mM) with magnesium supplementation (10.5 mM) to a DivCat: ASO ratio of 3:1. Note that increasing the volume injected to 100 µl of ASO (30 mg/ml; 4.22 mM) alone brings DivCat: ASO ratio 0.52:1 with a subsequent lower aA response; additionally, when 10.5 mM magnesium is supplemented to keep a DivCat: ASO ratio of 3:1 significant aA mitigation occurs. **(C)** Acute neuronal activation maximum response after 30 µl intrathecal administration of 3 mg dose of ASO C (100 mg/ml, 14.1 mM) alone or 100 µl IT of 3 mg dose of ASO C (30 mg/ml; 4.22 mM) alone or 100 µl IT of 3 mg dose of ASO C (30 mg/ml; 4.22 mM) with magnesium supplementation (10 mM) to a DivCat: ASO ratio of 2.9:1. Note how the experimental manipulation in B is reproduced in C when another ASO-inducing aA response is used. Data are expressed as mean ± s.d. (n≥ 4 females, body weight 218 ± 9 g; n≥ 4 males, body weight 287 ± 8 g). Each symbol represents an individual animal. The statistical significance was determined by the Kruskal-Wallis test. *P ≤ 0.05; ** P ≤ 0.01 *** P ≤ 0.001; **** P ≤ 0.0001 *vs* vehicle (aCSF) or ASO alone. Abbreviations: aCSF, artificial cerebrospinal fluid; IT, intrathecal; aA, acute neuronal activation.

### Increasing the divalent cation-to-oligonucleotide ratio mitigates the acute neuronal activation response in non-human primates

To determine if our mitigation approach translates to higher species, we examined the acute neuronal activation response in non-human primates (NHP) with or without magnesium supplementation in the dosing solutions formulated in aCSF. Cynomolgus monkeys were randomly assigned and maintained into three groups that were dosed in 4-week intervals, using five distinct dosing paradigms with ASO A (**Figure 7A, Supplementary Table 11**) First, one milliliter of ASO (70 mg/ml) was delivered by intrathecal administration formulated either in standard artificial cerebrospinal fluid or in aCSF supplemented with 21 mM magnesium resulting in a divalent cations-to-ASO ratio of 0.16:1 and 2.3:1, respectively. Consistent with our findings in rats, ASO A provoked acute neuronal activation responses in NHP, and the response was effectively mitigated with a 2.3:1 divalent cations-to-ASO ratio, maintaining the score similar to the score observed with aCSF vehicle (**Figure 7B**).

**Figure 7.**
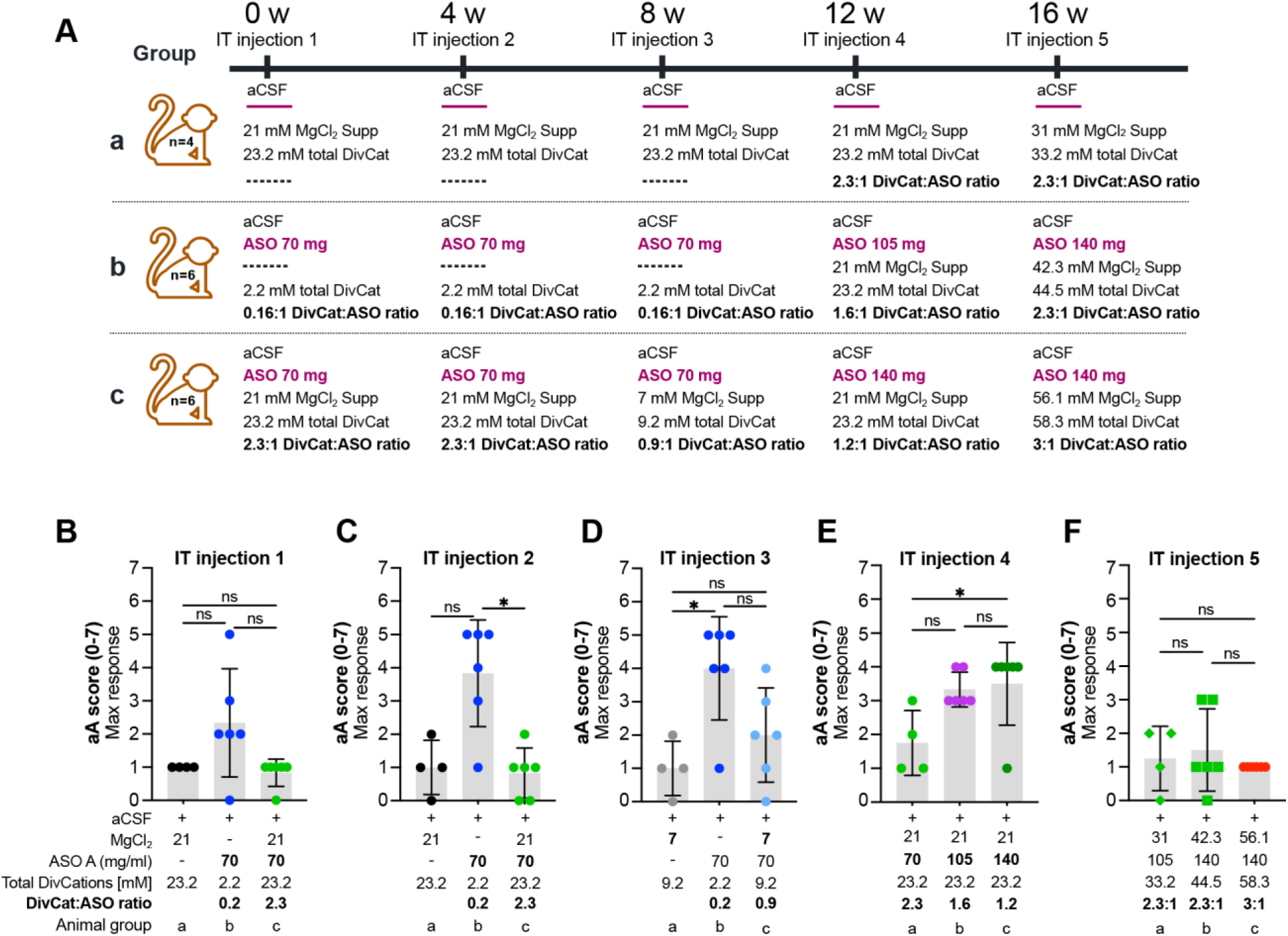
Increasing the divalent cation-to-oligo antisense ratio mitigates acute neuronal activation response in non-human primates. **(A)** Illustration of the dosing experimental design in NHP. **(B, C)** Acute neuronal activation (aA) maximum response in NHP after intrathecal administration of aCSF (black circles) or ASO A (70 mg/ml) alone (dark blue circles) or ASO A (70 mg/ml) with magnesium supplementation (21 mM) for a 2.3:1 DivCat: ASO ratio (green circles). The first (**B**) and second (**C**) doses were 4 weeks apart and the aA mitigation was reproduced when magnesium was supplemented for a 2.3:1 DivCat: ASO ratio. **(D)** Acute neuronal activation maximum response in NHP after intrathecal administration of aCSF with magnesium supplementation (7 mM) (light gray circles) or ASO A (70 mg/ml) alone (dark blue circles) or ASO A (70 mg/ml) with magnesium supplementation (7 mM) for a 0.9:1 DivCat: ASO ratio (light blue circles). Note how lowering magnesium supplementation and DivCat: ASO ratio mitigates aA to a lesser degree. **(E)** Acute neuronal activation maximum response in NHP after intrathecal administration of ASO A (70 mg/ml) with magnesium supplementation (21 mM) for a 2.3:1 DivCat: ASO ratio (green circles) or constant magnesium supplementation (21 mM) with increasing ASO A concentration to 105 mg/ml (purple circles) or 140 mg/ml (dark green circles) for a 1.6:1 or a 1.2:1 DivCat: ASO ratio, respectively. Note how increasing ASO concentration decreases the DivCat: ASO ratio, hence preventing aA mitigation. **(F)** Acute neuronal activation maximum response in NHP after intrathecal administration of increasing millimolar magnesium concentration supplementation and increasing ASO (mg/ml) concentration to maintain a DivCat: ASO ratio of 2.3:1 (green diamonds and green squares) or 3:1 (red circles). Data shows that when a DivCat: ASO ratio ≥ 2.3 is maintained, the aA mitigation is achieved. All five experiments were done in the same non-human primates with 4 weeks between each experiment, the animal groups used in each experiment were constant and are noted as a, b, and c. Each symbol represents an individual animal. Data are expressed as mean ± s.d. (n=4 females, body weight 2.6 ± 0.18 g and n=4 males, body weight 3.1 ± 0.59 kg). The significant difference was determined by Kruskal-Wallis test. *P ≤ 0.05; ** P ≤ 0.01 *** P ≤ 0.001; **** P ≤ 0.0001 *vs* the indicated groups. Abbreviations: aCSF, artificial cerebrospinal fluid; IT, intrathecal; aA, acute neuronal activation; DivCat, Divalent cations.

Four weeks later, this experiment was repeated in the same animals (IT injection 2). The acute neuronal activation response was still present after 70 mg of ASO A formulated in aCSF, and again mitigated by a 2.3:1 divalent cations-to-ASO ratio (Magnesium added) (**Figure 7C**).

In the third IT injection (8 weeks after the 1^st^ dose) the divalent cations-to-ASO ratio was modified from 2.3:1 to 0.9:1 by reducing the amount of magnesium added to the aCSF from 21 mM to 7 mM and keeping the ASO dose constant at 70 mg. The acute neuronal activation response was still present, and, as in our rat studies, a ratio of 0.9:1 divalent cations-to-ASO partially mitigated the acute neuronal activation responses (**Figure 7D**).

Next, twelve weeks after the 1^st^ dose (IT injection 4), we evaluated the response with increasing dose levels (70 mg, 105 mg, and 140 mg), and a constant magnesium supplementation at 21 mM in all groups. This produced a divalent cations-to-ASO ratio of 2.3:1. 1.6:1 and 1.2:1, respectively. Higher doses without the added magnesium were not tested. The 2.3:1 divalent cations-to-ASO ratio mitigated the acute neuronal activation responses, while the higher ASO doses with lower divalent cations-to-ASO ratios were not sufficient to fully prevent the response (**Figure 7E**).

Finally, in the last IT injection study (IT injection 5), our objective was to determine in NHP whether the acute neuronal activation responses were influenced by the ratio of cations-to-ASO in the dosing solution, rather than by the total ASO dose administered by maintaining a high molar ratio of divalent cations. One group of NHPs received 105 mg of ASO supplemented with 31 mM of magnesium, resulting in a cation-to-ASO ratio of 2.3:1. The second group received 140 mg of ASO with 42.3 mM of magnesium, maintaining the same 2.3:1 ratio. A third group received 140 mg of ASO supplemented with 56.1 mM of magnesium, increasing the ratio to 3:1. Acute neuronal activation response was mitigated when the divalent cations-to-ASO ratio was 2.3:1, moreover the scores were also comparable to vehicle levels with the ASO formulation that kept the divalent cations-to-ASO ratio at 3:1 (**Figure 7F**). Thus, with sufficient cation supplementation, acute neuronal activation responses were not observed with IT dosing up to 140 mg in NHPs.

Overall, the data supports the hypothesis that chelation of divalent cations by high ASO concentration causes the acute neuronal activation response and suggests that maintaining a molar ratio of divalent cations-to-ASO between 2.3:1 and 3:1 effectively mitigates the ASO-induced acute neuronal activation response in both NHP and rats.

### Divalent cations binding properties of oligonucleotides

To quantify the cation binding capabilities of ASO A, we dialyzed 10 mM ASO in buffers containing increasing concentrations of magnesium and calcium ions. As expected, both magnesium and calcium were bound by the ASO. With 10mM ASO, magnesium binding analysis revealed a binding saturation at 150.3 mM (**Figure 8B**), whereas calcium binding reached saturation at 60.8 mM (**Figure 8C**). Thus, fewer calcium ions (∼6 sodium ions replaced) were required to fully saturate the ASO compared to magnesium ions (∼15 sodium ions replaced). This is consistent with previous observations of siRNA metal ion binding (23), and our in vivo analysis that calcium can mitigate the acute activation effects at lower concentrations than magnesium (**Figure 3**).

**Figure 8.**
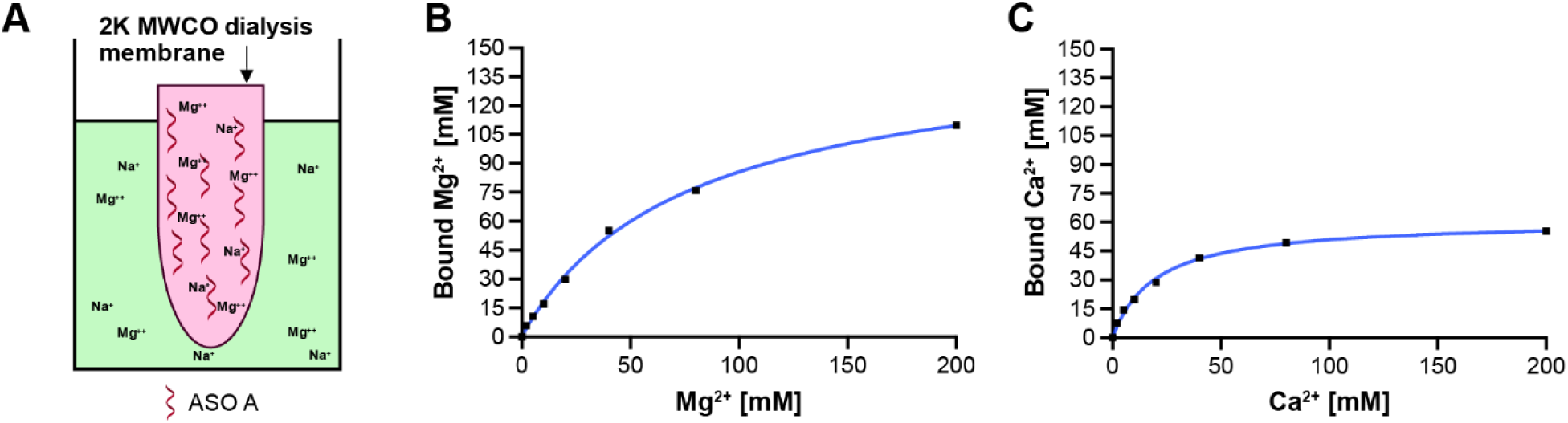
Oligo antisense binding properties of divalent cations. **(A)** Schematic of the microdialysis experiment showing 10 mM ASO A solutions prepared in buffers (10 mM sodium acetate [NaOAc] and 150 mM sodium chloride [NaCl], pH 7.2) with increasing concentrations of magnesium or calcium ions (0, 2, 5, 10, 40, 80, and 200 mM). **(B)** Plot of bound magnesium concentration of ASO A containing fraction against increasing magnesium concentration of dialysis buffer. **(C)** Plot of bound calcium concentration of ASO A containing fraction against increasing calcium concentration of dialysis buffer.

## DISCUSSION

We describe here an acute neuronal activation response in preclinical species following high-dose local administration of ON therapeutics into CSF. The acute neuronal activation response is transient, starting immediately after dosing or during recovery from anesthesia, with peak responses observed at 15-30 minutes post-dosing. Without intervention, the symptoms typically resolve after 2 hours, with no apparent sequelae. In mice, following ICV ON delivery the acute neuronal activation responses can present as hunching of the back, shivering/shaking, stiff and raised tail, hyperactivity, incoordination, hopping, and/or seizures. In rats following IT delivery of ON, the response presents as hunching of the back, shivering/shaking, muscle twitching, cramping, hyperactive/rapid movements, tail/hindlimb biting, stereotypic movements, defensive behaviors, vocalizations, and/or seizures. The acute neuronal activation responses also can occur in NHPs, and the response includes muscle tremors, twitching, salivation, self-biting, urinary incontinence, nystagmus, tonic or clonic movements, convulsions and rarely seizures with a similar time course of presentation as the rodents.

Our findings support a model where the acute neuronal activation response is caused by the high concentration of ON in the dosing solutions used in some preclinical studies which results in low divalent cations-to-ASO ratios leading to chelation of endogenous divalent cations by the ON in the CSF and possibly interstitial fluid of adjacent structures next to the site of injection. The timing of acute neuronal activation effects, immediately after dosing, appears to correlate with the Cmax of the ON and resolves as ASO distributes from the local injection site.

Total CSF volume varies greatly between species. Mouse CSF volume is ∼0.035-0.040 mL (40), rat CSF volume is ∼0.27-0.4 mL (41), NHP CSF volume is ∼7-15 mL, and human CSF volume is ∼ 96-150 mL (7, 40, 42). The dosing solution volume for safe intrathecal delivery is limited by the total intrathecal space, with typical delivery volumes of 5-10 ml in mice, < 0.1 mL in rats, 1.0-2.0 mL in NHP, and 15-20 mL in adult humans (14). As a result, the concentration of ASO in dosing solutions for preclinical studies is considerably higher than the concentrations for dosing in humans, at the same dose level. When formulated in aCSF with a fixed concentration of divalent cations (1.4 mM calcium and 0.8 mM magnesium), the preclinical formulations have much lower divalent cation-to-ON ratios. For example, Sprinraza has a divalent cation-to-ON ratio of 6.5 in the human formulation. The divalent cation-to-ASO ratio in NHP typically ranges from 0.2 to 1.5. Since the ASO formulation conditions in NHP do not reflect dosing in humans, where there are enough divalent cations in the formulation to prevent chelation of endogenous divalent cations in the CSF, this acute neuronal activation response is not expected to translate into the clinic at the dose levels and volumes currently being tested (43, 44).

The addition of magnesium to standard artificial cerebrospinal fluid (aCSF) used to prepare the ON preclinical formulation is a straightforward way to increase the divalent cations-to-ON ratio while maintaining the required dose and other factors (like volume) that affect ON pharmacology and distribution (45). One previously proposed strategy is to pre-saturate ONs with calcium before the delivery. However, this method can interfere with the solubility of some sequences (12). We also demonstrated here that calcium supplementation only provides a temporary reprieve from the acute neuronal activation effects and may cause other negative side effects or lead to precipitation of the ON. Intrathecal magnesium has been used broadly in clinical trials for pain management at extremely high doses and without lasting negative consequences (46–49). We propose an approach to empirically determine in vivo in rodents the amount of magnesium required for ON saturation to prevent the response. We establish here that this approach translates well to NHP when scaling based on the molar ratio of divalent cations-to-ON. This approach allows for empirical determination in rodents of the supplementation required to prevent the acute neuronal activation response, prior to expensive and time-consuming NHP studies.

Our work extends and clarifies previous observations in the field of acute responses in mice and rats following ON administration into the CSF. The previous neurological findings have been a mixture of inhibitory/sedation-like signs and excitatory/activation-like signs occurring between 30 minutes and 24 hours after dosing ASOs or siRNAs (9–12). Consistent with our findings, seizure-like behaviors have been observed in the first hour after dosing awake mice with siRNAs and were correlated with in vivo EMG/EEG high amplitude waves characteristic of seizures (23). Others have observed acute neuronal activation-like signs, such as hyperactivity, vocalization and seizures mostly occurring within minutes to 1 h after antisense oligonucleotide delivery and later describing sedation-like phenotypes within 2 to 4 h after dosing (9–12). After an extensive database search, we did not find a relationship between the acute neuronal activation and the acute sedation response and there is a clear separation in the timing of onset and resolution of the responses. Thus, it is likely that the ASOs used in the published assays were able to induce both acutely sedative and activating responses, and the paradigms and scoring systems utilized did not allow for a distinction between the unique responses, complicating the data interpretation. Here we propose a scoring system for rats, mice, and NHP that allows for accurate scoring of the acute neuronal activation response.

Our efforts clearly demonstrate that within a species, the likelihood and the severity of the response negatively correlate with the body weight of the animal. Moreover, the response of a given ON at a given dose is variable across animals, even in situations where body weight, gender, and genetics are controlled (i.e. inbred mice). Given the temporal dynamics and variability, it is likely that high local CSF concentrations of ON are needed to drive the response. CSF dynamics are linked to body-mass with a strong correlation between body mass index and cerebrospinal fluid pressure (50). CSF dynamics and flow are also altered with age (42, 51–53). Taken together, one plausible mechanism for the relationship between the acute neuronal activation response and body size, and the variability in the response, is that ON are more likely to be retained at high local CSF concentrations in the smaller animals, leading to a higher incidence and stronger response. Unfortunately, there are no existing methods to reliably measure these subtle changes in ON distribution and CSF flow immediately after dosing, and more work is needed to test this hypothesis. Understanding these variables is important in designing future studies evaluating the safety of ON therapeutics.

As the field of ON therapeutics expands, more research is required to apply these learnings to new chemistries and ON therapeutic designs and to continue to understand the underlying mechanisms of ON toxicities, so that only the most effective and safest ON are provided to patients.

## Supporting information

Bravo Hernandez_aA Supplement

## ACKNOWLEDGEMENTS

The authors thank Donna Sipe and the vivarium staff, oligo synthesis group, PCR core group, histology core staff, Raul Alonzo, and the creative services team at Ionis Pharmaceuticals Inc., for their technical support.

## AUTHOR CONTRIBUTIONS

Mariana Bravo Hernandez: conceptualization, data curation, formal analysis, investigation, methodology, project administration, visualization, writing-original draft. Curt Mazur: conceptualization, data curation, formal analysis, investigation, methodology, writing-review & editing. Hao Chen: Conceptualization, data curation, formal analysis, methodology, writing-review & editing. Linda Fradkin: data curation, methodology, resources, writing-review & editing. Justin Searcy: methodology, resources, writing-review & editing. Sebastien Burel: Data curation, Formal analysis, Investigation, methodology, software, visualization, writing-review & editing. Mackenzie Kelly: investigation Dona Bruening: investigation. Jacqueline G. O’Rourke: writing-review & editing. Yuhang Cai: resources. Jonathon Nguyen: investigation, Lisa Berman-Booty: data curation, investigation, visualization, writing-review & editing. Lendell Cummins: data curation, formal analysis, investigation, visualization. Hans Gaus: formal analysis, investigation, methodology, validation, visualization writing-review & editing. Berit Powers: writing-original draft. Hien Zhao: resources, writing-review & editing. Paymaan Jafar-Nejad: supervision, validation, writing-review & editing. Scott Henry: supervision, validation, writing-review & editing, Eric Swayze. supervision, validation, writing-review & editing. Holly B. Kordasiewicz: conceptualization, resources, supervision, validation, writing-original draft.

## SUPPLEMENTARY DATA

Supplementary Data are available online.

## CONFLICT OF INTEREST

MBH, HC, LF, JS, SB, MK, DB, JGO, YC, JN, LBB, LC, HG, BP, HZ, PJN, SH, ES, and HBK are employees and shareholders of IONIS. CM is a former employee and shareholder of IONIS.

## FUNDING

Studies were funded by IONIS. Funding for open access charge: IONIS.

