## Supplementary material for "Transient Acute Neuronal Activation Response Caused by High Concentrations of Oligonucleotides in the Cerebral Spinal Fluid": Bravo Hernandez_aA Supplement

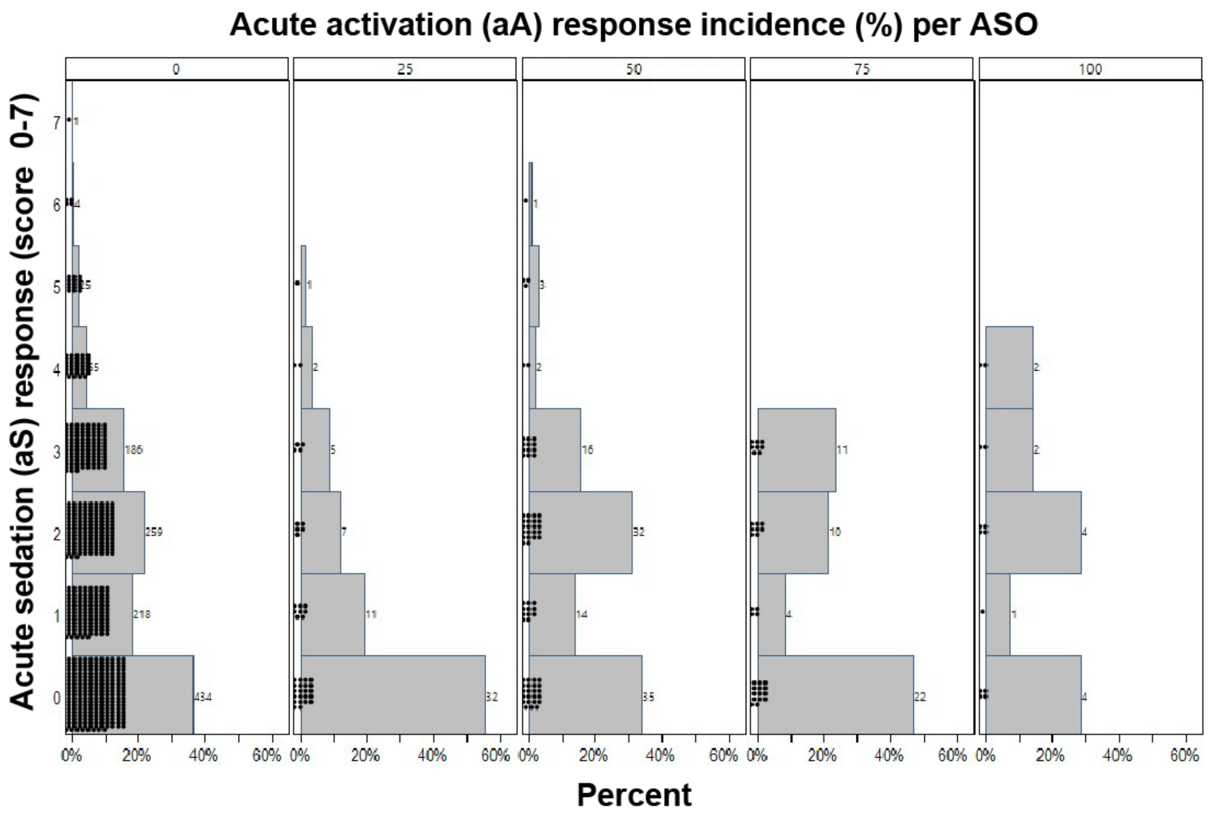


**Supplementary Figure 1. The acute sedation (aS) response does not have an associated increased incidence of acute neuronal activation (aA) response.** Comparing the severity of acute sedation using the acute sedation scale (score 0-7) at 3 h after ASO administration (3 mg dose) to the incidence of acute neuronal activation (aA) response. Each panel represents a level of aA incidence from 0 to 100% in 25% increments. The histogram within each panel is expressed as the percentage of ASO with a given level of acute sedation score (average score per ASO). Each dot represents a unique ASO tested in independent studies with n≥ 4 rats in each experimental group. The figure suggests there is no connection between the occurrence of acute sedation and the presence of acute neuronal activation response.

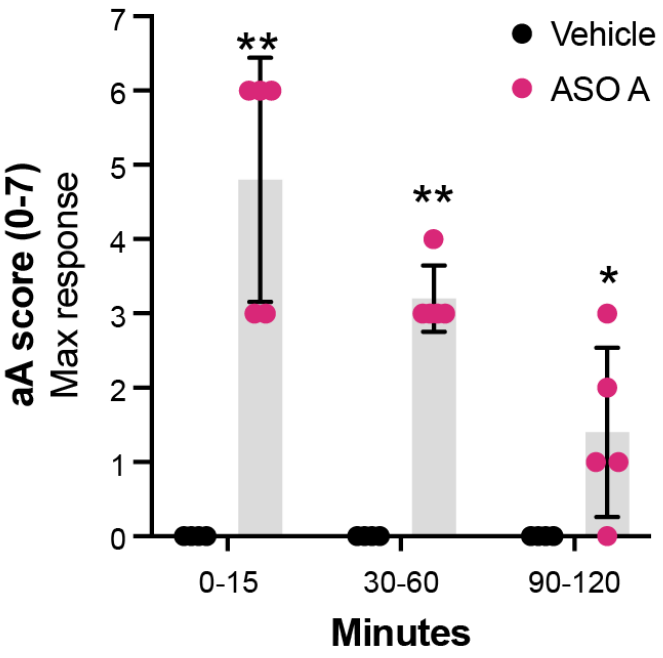


**Supplementary Figure 2. Time-course of acute neuronal activation response in female rats.** Acute neuronal activation maximum response in female rats (body weight 204 ± 4 g) after a single IT administration of 3 mg ASO A in 30 µl (100 mg/ml) (pink circles, n= 5) or 30 µl of artificial cerebrospinal fluid (Vehicle, black circles, n= 4). The aA scores for each group (ASO A or Vehicle), were divided into three-time blocks (0-15 minutes, 30-60 minutes, and 90-120 minutes. Data are expressed as mean ± s.d. The statistical significance was determined by the Mann-Whitney test. *P ≤ 0.05; ** P ≤ 0.01 *** P ≤ 0.001; **** P ≤ 0.0001 *vs* vehicle (aCSF). Abbreviations: IT; intrathecal; aA= acute neuronal activation; aCSF=artificial cerebrospinal fluid.

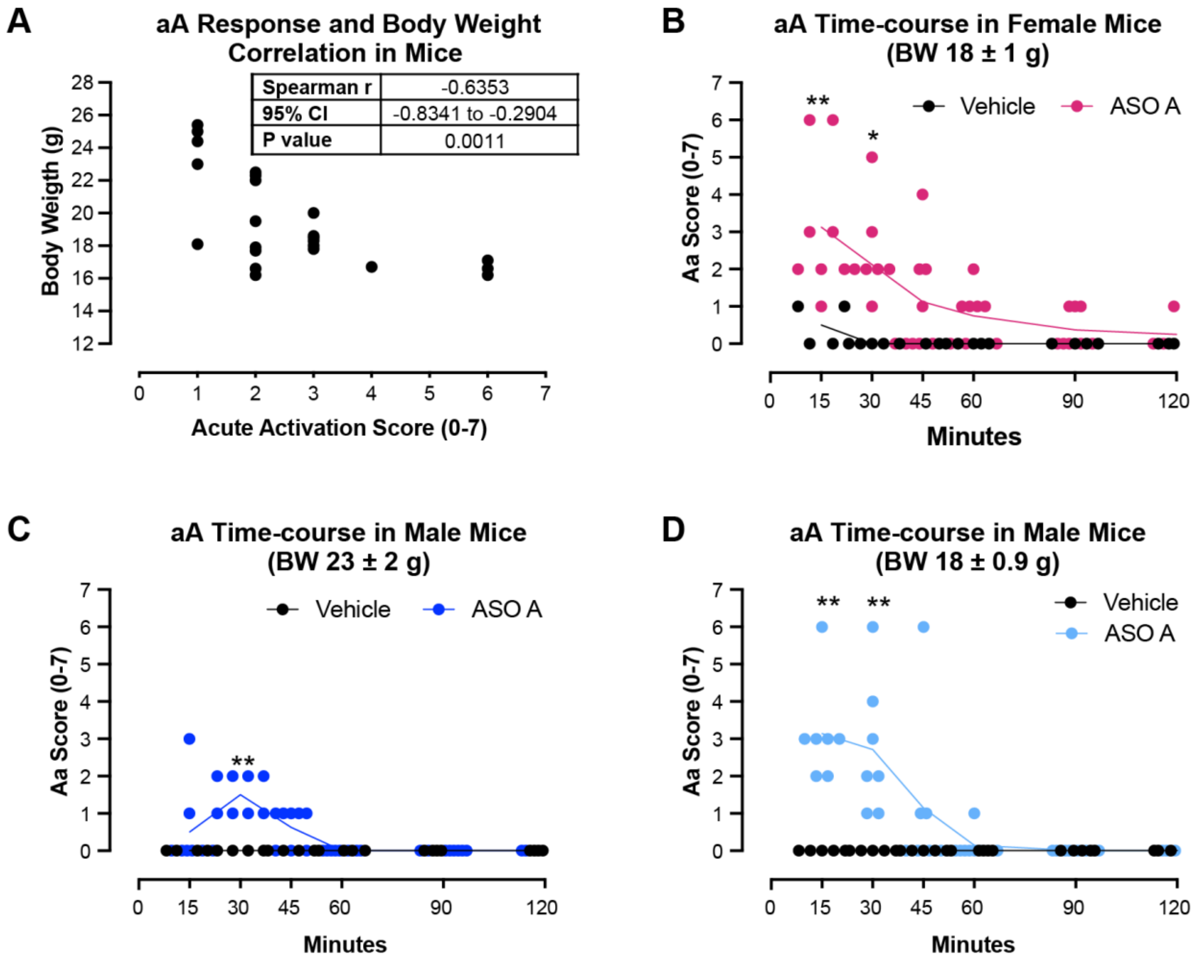


**Supplementary Figure 3. Body-weight dependent, transient acute neuronal activation response after oligonucleotide injection in mice. (A)** Correlation graph of acute neuronal activation (aA) score versus mouse body weight; each dot represents a unique animal. Note how animals with lower body weight had a higher aAtox score. **(B-D)** Time-course of aA score after single intracerebroventricular (ICV) administration of 1000 µg ASO A in 10 µl (100mg/ml, 14.1 mM) (n≥ 7) or 10 µl of vehicle (aCSF; n=4, black circles) in females with BW ~18 g (**B**), Males with BW ~ 23 g (**C**) or Males with BW ~18 g (**D**). Each symbol represents an individual animal. The data is expressed as the mean ± s.d. The statistical significance was determined as follows: Spearman r correlation and linear regression (A); Multiple Mann-Whitney test (B-D). *P ≤ 0.05; ** P ≤ 0.01 *** P ≤ 0.001; **** P ≤ 0.0001 *vs* vehicle (aCSF). Abbreviations: ICV=intracerebroventricular; aA= acute neuronal activation; aCSF=artificial cerebrospinal fluid.

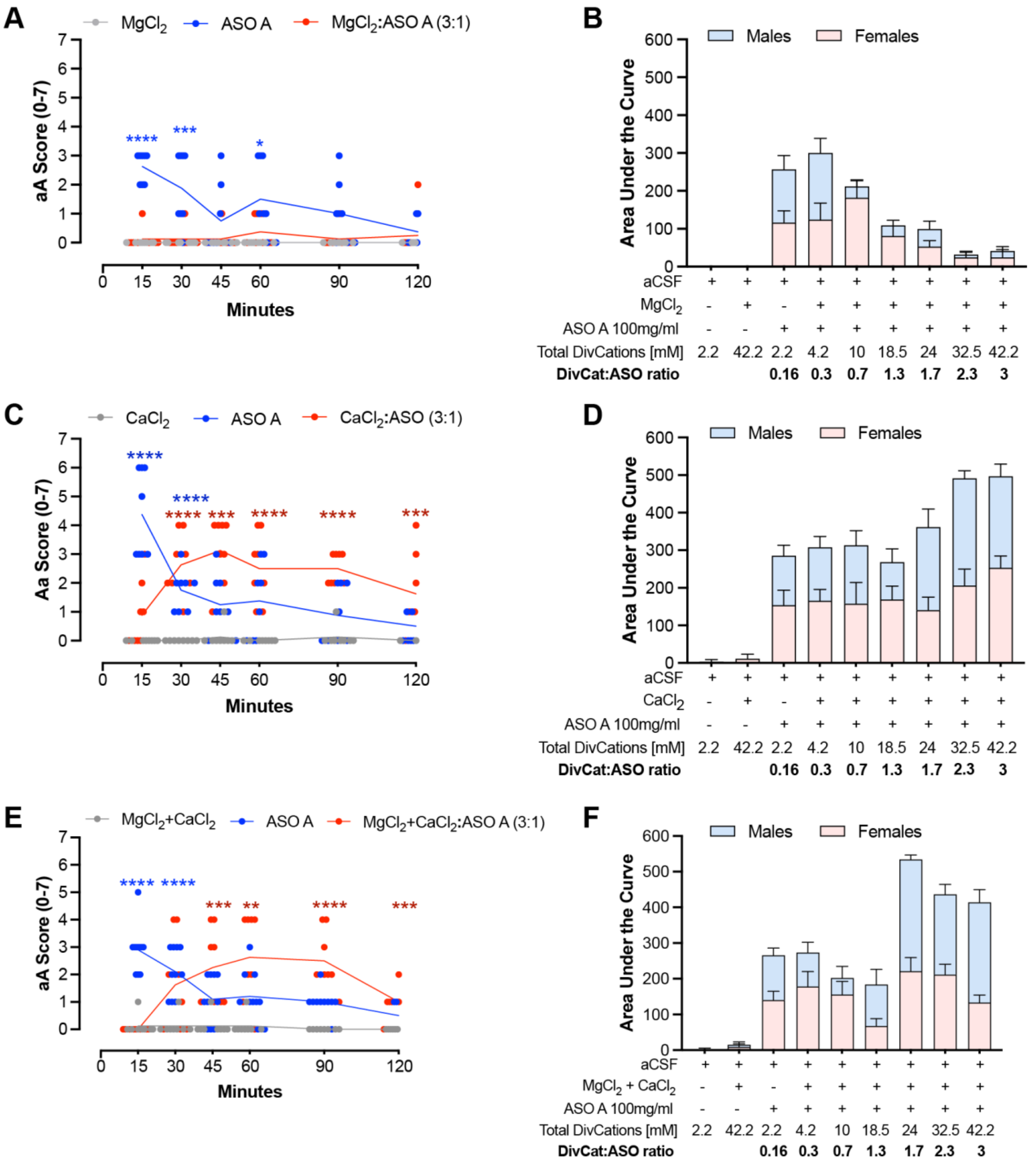


**Supplementary Figure 4. Time course of ASO-induced acute neuronal activation response modifying the divalent cation-to-oligo antisense ratio with magnesium or calcium supplementation or the combination of both. (A-B)** Time course (**A**) and area under the curve (**B**; females in pink, males in blue) of acute neuronal activation (aA) response in rats after intrathecal 3 mg dose administration of ASO A (100 mg/ml, 14.1 mM) formulated in artificial cerebrospinal fluid (aCSF) with or without increasing millimolar concentrations of magnesium supplementation, resulting in six different DivCat: ASO ratio from 0.3:1 to 3:1 (n≥ 4 females, BW=

225 ± 8 g and n≥ 4 males, BW= 286 ± 11 g). The data shows a significant reduction of aA response in a ratio-dependent manner over the 2 hours. **(C-D)** Time course (**C**) and area under the curve (**D**; females in pink, males in blue) of acute neuronal activation response in rats after intrathecal 3 mg dose administration of ASO A (100 mg/ml, 14.1 mM) formulated in artificial cerebrospinal fluid (aCSF) with or without increasing millimolar concentrations of calcium supplementation, resulting in six different DivCat: ASO ratio from 0.3:1 to 3:1 (n≥ 4 females, BW= 220 ± 10 g and n≥ 4 males, BW= 285 ± 10 g). The data shows a clear, sustained, and significant rebound acute activation response effect from the 30 min over the 2 hours. **(E-F)** Time course (**E**) and area under the curve (**F**; females in pink, males in blue) of acute neuronal activation response in rats after intrathecal 3 mg dose administration of ASO A (100 mg/ml, 14.1 mM) formulated in artificial cerebrospinal fluid (aCSF) with or without increasing millimolar concentrations of magnesium plus calcium supplementation (kept at their physiological ratio), resulting in six different DivCat: ASO ratio from 0.3:1 to 3:1 (n≥ 4 females, BW= 224 ± 8 g and n≥ 4 males, BW= 294 ± 8 g). The data shows a clear, sustained, and significant rebound acute activation response effect from 30 min to 2 hours with a higher AUC than the AUC of ASO alone at the highest ratios tested. Each symbol represents an individual animal. Data are expressed as the time course of the aA response (A, C, E) or the area under the curve (mean aA score within 120 min) (B, D, F). The statistical significance was determined by the Multiple Mann-Whitney test (A, C, E), and the area under the curve (AUC) was calculated using the trapezoid rule (B, D, F). P ≤ 0.05; ** P ≤ 0.01 *** P ≤ 0.001; **** P ≤ 0.0001 *vs* aCSF + 42.2 mM magnesium, calcium or magnesium+calcium respectively. Abbreviations: aA= acute neuronal activation; AUC= area under the curve; aCSF=artificial cerebrospinal fluid.

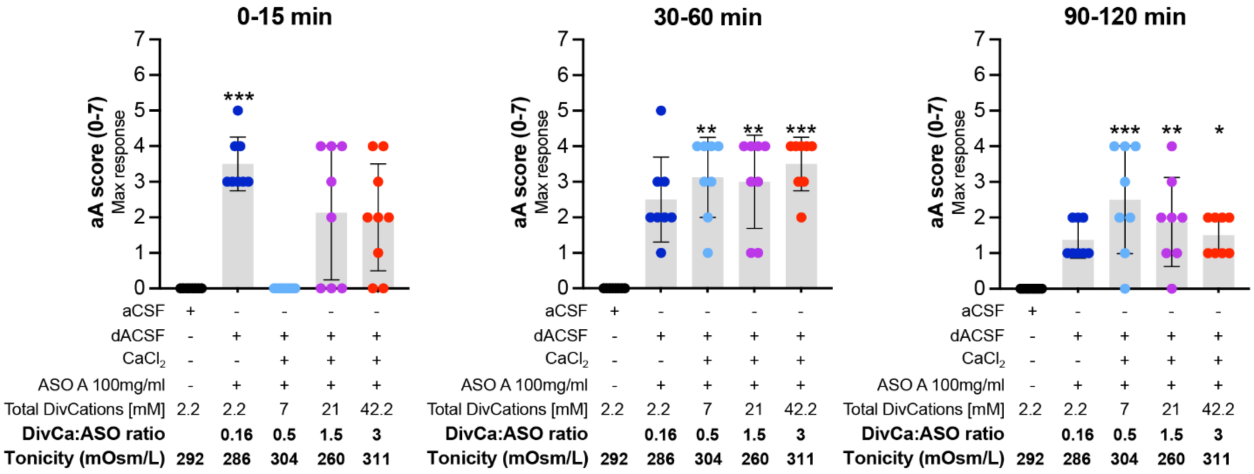


**Supplementary Figure 5. Effect of increasing the divalent cation-to-oligo antisense ratio by calcium supplementation under isotonic conditions.** Acute neuronal activation (aA) maximum response plotted in three-time blocks; 0-15 minutes (left); 30-60 minutes (middle); and 90-120 minutes (right) after intrathecal 3 mg dose administration of ASO A (100 mg/ml, 14.1 mM) formulated in the sodium-deficient artificial cerebrospinal fluid (dACSF) with or without increasing millimolar concentrations of calcium supplementation, resulting in six different DivCat: ASO ratio from 0.5:1 to 3:1 (n≥ 4 females, BW= 225 ± 2 g and n≥ 4 males, BW= 300 ± 5 g). The data shows a significant reduction of aA response with a 0.5:1 DivCat: ASO, but not 1.5:1 or 3:1 at 0-15 min time point. A clear, sustained, and significant rebound acute activation effect was observed at all tested ratios from 30 min up to 2 hours. The dACSF was not tested alone as tonicity is under physiological levels. Each symbol represents an individual animal. The data is expressed as the mean ± s.d. The statistical significance was determined by the Kruskal-Wallis test. *P ≤ 0.05; ** P ≤ 0.01 *** P ≤ 0.001; **** P ≤ 0.0001 *vs* vehicle (dACSF). Abbreviations: aA= acute neuronal activation; aCSF=artificial cerebrospinal fluid; dACSF= sodium deficient artificial cerebrospinal fluid (isotonic)**.**

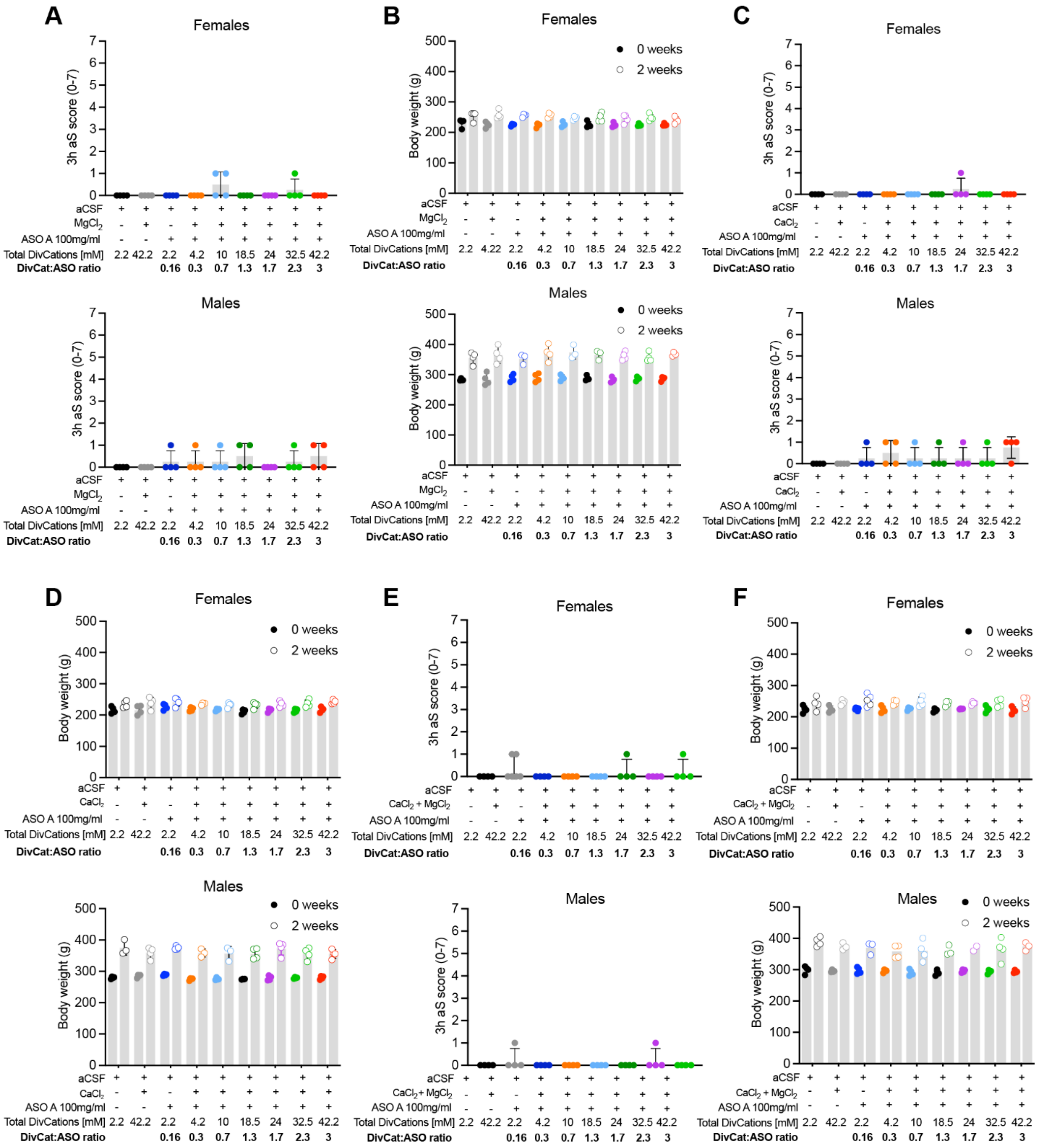

**Supplementary Figure 6. Modifying the divalent cation-to-oligo antisense ratio with magnesium or calcium supplementation or the combination of both do not induce acute sedation phenotype, nor affect body weight gain. (A)** Acute sedation response in female (top) and male (bottom) rats 3 hours after intrathecal administration of 3 mg/ 30 µl of ASO A (100mg/ml, 14.1 mM), with or without varying millimolar concentrations of magnesium supplementation formulated in aCSF, and **(B)** body weight gain in both sexes (top females, bottom males) after 2 weeks of intrathecal dosing of 3 mg/ 30 µl of ASO A (100 mg/ml), also with or without magnesium supplementation resulting in six different DivCat: ASO ratios, ranging from 0.3:1 to 3:1 (n≥ 4 females, BW= 225 ± 8 g and n≥ 4 males, BW= 286 ± 11 g). **(C)** Acute sedation response in female (top) and male (bottom) rats 3 hours after intrathecal administration of 3 mg/ 30 µl of ASO A, with or without varying millimolar concentrations of calcium supplementation formulated in aCSF, and **(D)** body weight gain in both sexes (top females, bottom males) after 2 weeks of intrathecal dosing (3 mg ASO A at 100 mg/ml), also with or without calcium supplementation resulting in six different DivCat: ASO ratios, ranging from 0.3:1 to 3:1 (n≥ 4 females, BW= 220 ± 10 g and n≥ 4 males, BW= 285 ± 10 g). (**E)** Acute sedation response in female (top) and male (bottom) rats 3 hours after intrathecal administration of 3 mg/30 µl of ASO A (100mg/ml, 14.1 mM), with or without varying millimolar concentrations of magnesium plus calcium supplementation (kept at their physiological ratio) formulated in aCSF, and **(F)** body weight gain in both sexes (top females, bottom males) after 2 weeks of intrathecal dosing (3 mg ASO A at 100 mg/ml), also with or without magnesium plus calcium supplementation resulting in six different DivCat: ASO ratios, ranging from 0.3:1 to 3:1 (n≥ 4 females, BW= 224 ± 8 g and n≥ 4 males, BW= 294 ± 8 g). Body weight gain was not affected in any experimental group, and no relevant acute sedation response was observed. Each point represents an individual animal. Data are expressed as mean ± s.d. The statistical significance was determined by the Kruskal-Wallis test. *P ≤ 0.05; ** P ≤ 0.01 *** P ≤ 0.001; **** P ≤ 0.0001 *vs* vehicle (aCSF). Abbreviations: aCSF; artificial cerebrospinal fluid; aA, acute neuronal activation.

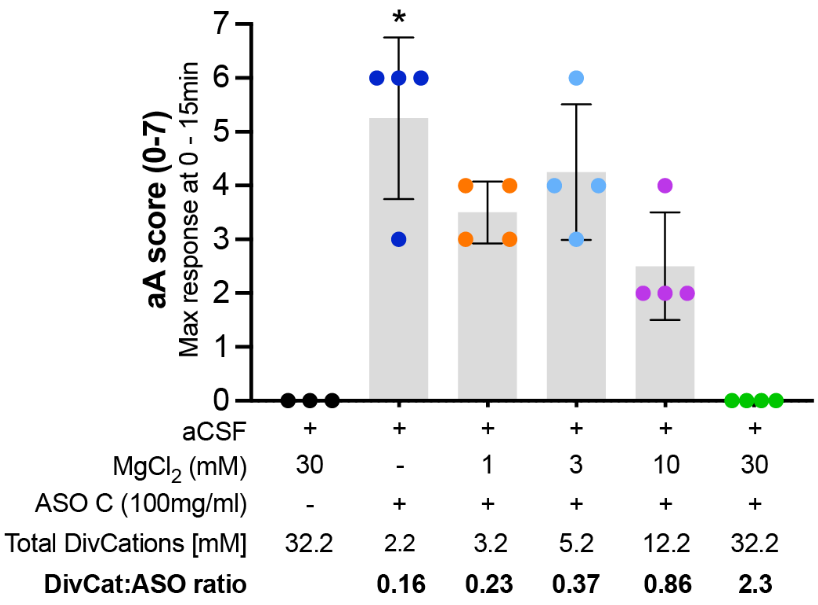


**Supplementary Figure 7. ASO C-induced-acute neuronal activation is mitigated by modifying the divalent cation-to-oligo antisense ratio with magnesium supplementation.** Acute neuronal activation (aA) maximum response 0-15 min after intrathecal 3 mg dose administration of ASO C (100 mg/ml,14.1 mM) with or without increasing millimolar concentrations of magnesium supplementation formulated in aCSF, resulting in four different DivCat: ASO ratio from 0.23:1 to 2.3:1 (n=4 male rats, BW= 263 ± 8 g). The data shows a significant reduction of aA response in a ratio-dependent manner. Each point represents an individual animal. Data are expressed as mean ± s.d. The statistical significance was determined by the Kruskal-Wallis test. *P ≤ 0.05; ** P ≤ 0.01 *** P ≤ 0.001; **** P ≤ 0.0001 *vs* vehicle (aCSF). Abbreviations: aCSF; artificial cerebrospinal fluid; aA, acute neuronal activation.

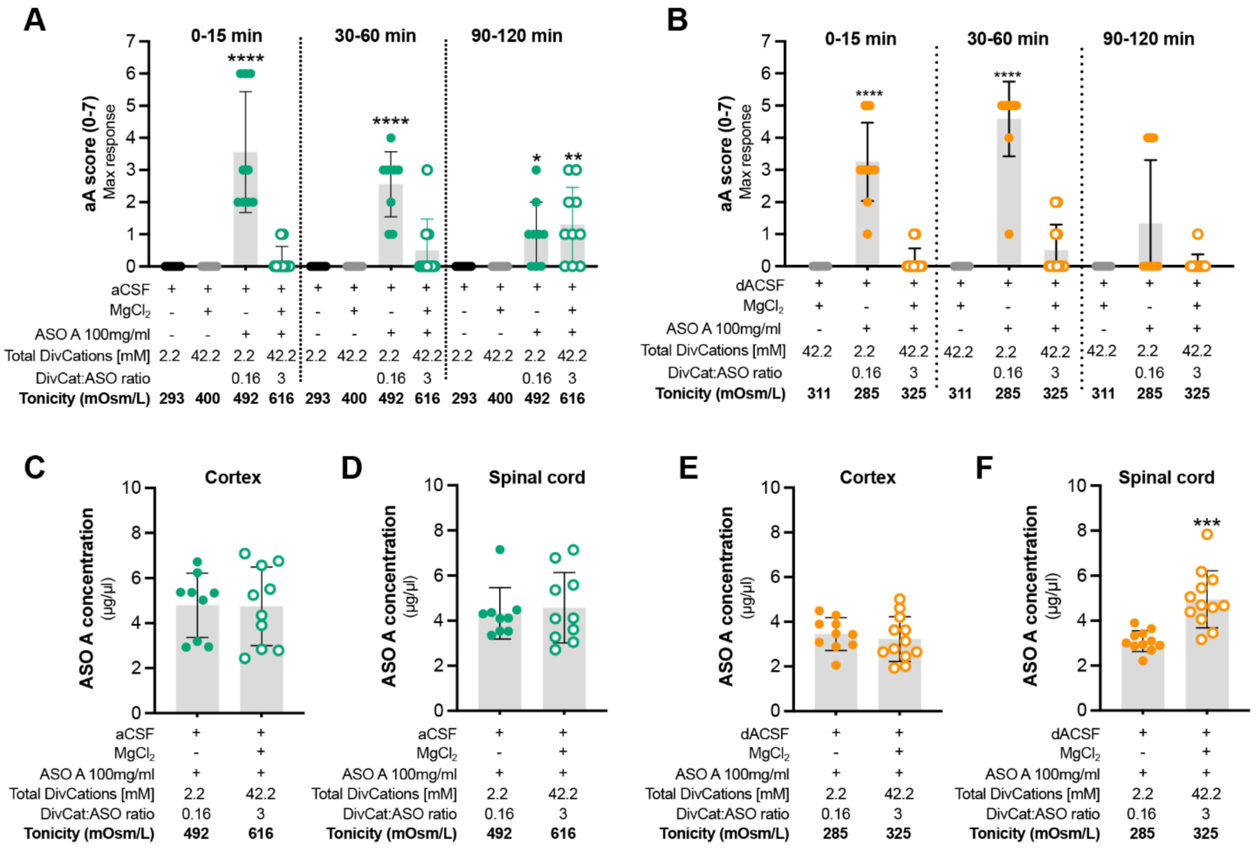


**Supplementary Figure 8. Modifying the divalent cation-to-oligo antisense ratio with magnesium supplementation mitigates ASO-induced acute neuronal activation response under hypertonic or isotonic conditions. (A)** Acute neuronal activation maximum response plotted in three-time blocks; 0-15 minutes (left); 30-60 minutes (middle); and 90-120 minutes (right) after intrathecal 3 mg dose administration of ASO A (100 mg/ml,14.1 mM) with or without 40 mM of magnesium supplementation for a 3:1 DivCat: ASO ratio formulated in artificial cerebrospinal fluid (aCSF). Note that formulation in aCSF with ASO and magnesium increases tonicity from 492 to 615 mOsm/L. (n≥ 4 females, BW= 204 ± 4 g and n≥ 4 males, BW= 261 ± 5 g). **(B)** Acute neuronal activation response maximum response plotted in three-time blocks; 0-15 minutes (left); 30-60 minutes (middle); and 90-120 minutes (right) after intrathecal 3 mg dose administration of ASO A (100 mg/ml,14.1 mM) with or without 40 mM of magnesium supplementation for a 3:1 DivCat: ASO ratio formulated in tonicity controlled (isotonic) artificial cerebrospinal fluid (sodium deficient aCSF, dACSF). Note that formulation in dACSF with ASO and magnesium keeps tonicity closer to physiological levels (325 mOSm/L) but does not affect the acute neuronal activation mitigation effect (n≥ 4 females, BW= 201 ± 3 g and n≥ 4 males, BW= 271 ± 2 g). **(C-D)** ASO tissue accumulation in the cortex (**C**) or spinal cord (**D**) of rats treated with intrathecal 3 mg dose of ASO A (100 mg/ml, 14.1 mM) with or without magnesium supplementation in standard artificial cerebrospinal fluid (aCSF). **(E-F)** ASO tissue accumulation in the cortex (**E**) or spinal cord (**F**) of rats treated with intrathecal 3 mg dose of ASO A (100 mg/ml, 14.1 mM) with or without magnesium supplementation formulated in tonicity-controlled artificial cerebrospinal fluid (dACSF). Each point represents an individual animal. Data are expressed as mean ± s.d. The statistical significance was determined as follows: Kruskal-Wallis test *vs* vehicle (A, aCSF; B, dACSF + magnesium) and Unpaired t-test (C-F). *$P \leq 0.05$; ** $P \leq 0.01$ *** $P \leq 0.001$; **** $P \leq 0.0001$. Abbreviations: aCSF= artificial cerebrospinal fluid; dACSF= sodium deficient aCSF; aA, acute neuronal activation.

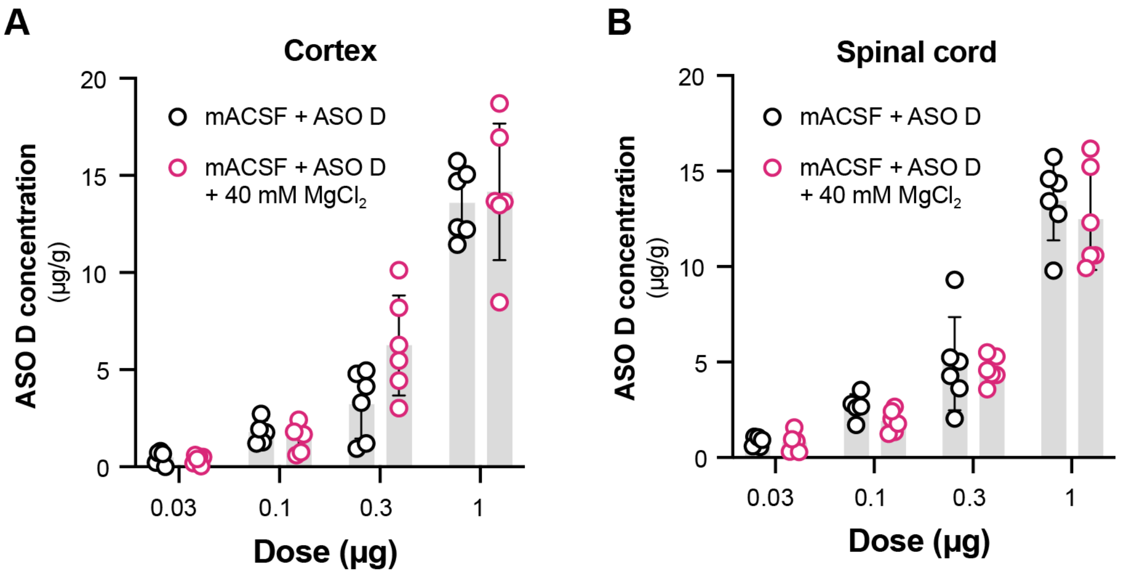


**Supplementary Figure 9. Dose-dependent ASO tissue concentration is observed with or without magnesium supplementation in rats. (A-B).** ASO tissue concentration plotted against increasing doses of ASO D (100 mg/ml) in the cortex (**A**) and spinal cord (**B**) after 2 weeks of intrathecal administration with or without magnesium supplementation (40 mM) formulated in a modified aCSF (mACSF; magnesium and calcium removed) in rats. Each point represents an individual animal (n=6 males, BW= 305 ± 9 g). Data are expressed as mean ± s.d. The significant difference was determined by the Unpaired t-test . *P ≤ 0.05; ** P ≤ 0.01 *** P ≤ 0.001; **** P ≤ 0.0001. No significant differences were found. Abbreviations: mACSF= modified artificial cerebrospinal fluid.

Table supplementary 1. ASO sequences

| ID | Simple sequence |
|---|---|
| ASO A | GCoAoCoCATATATATCTCoAoGAA |
| ASO B | AGoCoCoAATATTTATAGGoToGCT |
| ASO C | GToToGoGCCTTATCATACoCoTAA |
| ASO D | GCoCoAoGoGCTGGTTATGAoCoTCA |
| ASO E | GToAoToCAGTCATTATCToCoATC |
| ASO F | AToToCoTCTTCATCTATCoAoCCC |
| ASO G | TToGoCoTTCACAACTCCCoAoCCT |
| ASO H | TGoToCoCCTAACATTTTCoCoTTA |
| ASO I | CToGoCoCAGACATATATCoAoTAA |
| ASO J | AAoToGoCTTTCATAACTToCoCAC |
| ASO K | GCoToCoATTTATTCTCAAoGoTAC |
| ASO L | TAoGoGoAGTTAACTAGAAoCoTCA |
| ASO M | CCoGoGoAAGTCTTCTCTAoAoGTC |
| ASO N | TCoToCoAGCTCCTTCCTAoGoGGC |
| ASO O | GCoAoAoATTGATTACCTCoAoGTC |
| ASO P | ACoToAoGCGGTTATCTTCoCoAAC |
| ASO Q | GToAoToGCTACATTCAAAoToCTA |
| siRNA A | TGAoAoCoUoUoGoUoAoGoGoUoUoGoGoAoUoUoUoUCG<br>AAAoAoUoCoCoAoAoCoCoUoAoCoAoAoGoUoUCA |

Orange= 2'-O-methoxyethylribose; Black=DNA; PO= o; Yellow= 2'-O-MOE-(E)-5'-vinylphosphonate ;Green= 2'-Fluororibose; Red= 2'-O-methylribose; Gray= 2'-O-hexadecylribose . *All linkages are PS unless noted

**Table supplementary 2.** Acute sedation (aS) scoring following IT administration in rodents

| 3h aS score | Rodent IT injection acute sedation scale |
|---|---|
| **0** | Bright, alert, responsive |
| **1** | No tone/movement in tail |
| **2** | Weak posterior posture |
| **3** | Hind limbs don’t support weight, but can still move |
| **4** | Hind paws don’t move – full hindlimb paralysis |
| **5** | Weak anterior posture |
| **6** | Fore paws don’t move, but animal is still breathing |
| **7** | Death |

Rats are observed 3 hours after dosing and a score is assigned

**Table supplementary 3.** Formulation description of ASO A with or without increasing concentrations of magnesium supplementation.

| Treatment group | ASO Conc. (mg/ml) | ASO Molarity (mM) | $Ca^{++}$ Molarity (mM) | $Mg^{++}$ Molarity (mM) | $MgCl_2$ supplemented Molarity (mM) | DivCat: ASO ratio |
|---|---|---|---|---|---|---|
| aCSF | 0 | 0 | 1.4 | 0.8 | 0 | **0** |
| aCSF+ 32.5 mM total DivCat (Mg supp) | 0 | 0 | 1.4 | 0.8 | 30.3 | **0** |
| aCSF+ 42.2 mM total DivCat (Mg supp) | 0 | 0 | 1.4 | 0.8 | 40 | **0** |
| aCSF+ ASO 100 mg | 100 | 14.1 | 1.4 | 0.8 | 0 | **0.16:1** |
| aCSF+ 4.2 mM total DivCat (Mg supp)+ ASO 100 mg | 100 | 14.1 | 1.4 | 0.8 | 2 | **0.3:1** |
| aCSF+ 10 mM total DivCat (Mg supp)+ ASO 100 mg | 100 | 14.1 | 1.4 | 0.8 | 7.8 | **0.7:1** |
| aCSF+ 18.5 mM total DivCat (Mg supp)+ ASO 100 mg | 100 | 14.1 | 1.4 | 0.8 | 16.3 | **1.3:1** |
| aCSF+ 24 mM total DivCat (Mg supp)+ ASO 100 mg | 100 | 14.1 | 1.4 | 0.8 | 21.8 | **1.7:1** |
| aCSF+ 32.5 mM total DivCat (Mg supp)+ ASO 100 mg | 100 | 14.1 | 1.4 | 0.8 | 30.3 | **2.3:1** |
| aCSF+ 42.2 mM total DivCat (Mg supp)+ ASO 100 mg | 100 | 14.1 | 1.4 | 0.8 | 40 | **3:1** |

All groups were dosed at 30 mg/ 30µl. Abbreviations: DivCat= divalent cations; Mg supp= magnesium supplemented; aCSF= standard artificial cerebrospinal fluid

**Table supplementary 4.** Formulation description of ASO A with or without increasing concentrations of calcium supplementation.

| Treatment group | ASO Conc. (mg/ml) | ASO Molarity (mM) | $Ca^{++}$ Molarity (mM) | $Mg^{++}$ Molarity (mM) | $MgCl_2$ supplemented Molarity (mM) | DivCat: ASO ratio |
|---|---|---|---|---|---|---|
| aCSF | 0 | 0 | 1.4 | 0.8 | 0 | **0** |
| aCSF+ 42.2 mM total DivCat (Ca supp) | 0 | 0 | 1.4 | 0.8 | 40 | **0** |
| aCSF+ ASO 100 mg | 100 | 14.1 | 1.4 | 0.8 | 0 | **0.16:1** |
| aCSF+ 4.2 mM total DivCat (Ca supp)+ ASO 100 mg | 100 | 14.1 | 1.4 | 0.8 | 2 | **0.3:1** |
| aCSF+ 10 mM total DivCat (Ca supp)+ ASO 100 mg | 100 | 14.1 | 1.4 | 0.8 | 7.8 | **0.7:1** |
| aCSF+ 18.5 mM total DivCat (Ca supp)+ ASO 100 mg | 100 | 14.1 | 1.4 | 0.8 | 16.3 | **1.3:1** |
| aCSF+ 24 mM total DivCat (Ca supp)+ ASO 100 mg | 100 | 14.1 | 1.4 | 0.8 | 21.8 | **1.7:1** |
| aCSF+ 32.5 mM total DivCat (Ca supp)+ ASO 100 mg | 100 | 14.1 | 1.4 | 0.8 | 30.3 | **2.3:1** |
| aCSF+ 42.2 mM total DivCat (Ca supp)+ ASO 100 mg | 100 | 14.1 | 1.4 | 0.8 | 40 | **3:1** |

All groups were dosed at 30 mg/ 30µl. Abbreviations: DivCat= divalent cations; Ca supp= calcium supplemented; aCSF= standard artificial cerebrospinal fluid

**Table supplementary 5.** Formulation description of ASO A with or without increasing concentrations of calcium + magnesium supplementation.

| Treatment group | ASO Conc. (mg/ml) | ASO Molarity (mM) | $Ca^{++}$ Molarity (mM) | $Mg^{++}$ Molarity (mM) | $CaCl_2$ suppl Molarity (mM) | $MgCl_2$ suppl Molarity (mM) | DivCat: ASO ratio |
|---|---|---|---|---|---|---|---|
| aCSF | 0 | 0 | 1.4 | 0.8 | 0 | 0 | **0** |
| aCSF+32.5 mM total DivCat (Mg+Ca supp) | 0 | 0 | 1.4 | 0.8 | 19.3 | 11.0 | **0** |
| aCSF+42.2 mM total DivCat (Mg+Ca supp) | 0 | 0 | 1.4 | 0.8 | 25.5 | 14.6 | **0** |
| aCSF+ASO 100 mg | 100 | 14.1 | 1.4 | 0.8 | 0 | 0 | **0.16:1** |
| aCSF+4.2 mM total DivCat (Mg+Ca supp)+ASO 100 mg | 100 | 14.1 | 1.4 | 0.8 | 1.29 | 0.74 | **0.3:1** |
| aCSF+10 mM total DivCat (Mg+Ca supp)+ASO 100 mg | 100 | 14.1 | 1.4 | 0.8 | 4.96 | 2.84 | **0.7:1** |
| aCSF+18.5 mM total DivCat (Mg+Ca supp)+ASO 100 mg | 100 | 14.1 | 1.4 | 0.8 | 10.4 | 5.9 | **1.3:1** |
| aCSF+24 mM total DivCat (Mg+Ca supp)+ASO 100 mg | 100 | 14.1 | 1.4 | 0.8 | 15.8 | 9.0 | **1.7:1** |
| aCSF+32.5 mM total DivCat (Mg+Ca supp)+ASO 100 mg | 100 | 14.1 | 1.4 | 0.8 | 19.3 | 11.0 | **2.3:1** |
| aCSF+42.2 mM total DivCat (Mg+Ca supp)+ASO 100 mg | 100 | 14.1 | 1.4 | 0.8 | 25.5 | 14.6 | **3:1** |

All groups were dosed at 30 mg/ 30µl. Abbreviations: DivCat= divalent cations; Mg+Ca supp= Magnesium plus calcium supplemented; aCSF= standard artificial cerebrospinal fluid

**Table supplementary 6.** Formulation description of ASO A with or without increasing concentrations of calcium supplementation formulated in dACSF (isotonic).

| Treatment group | ASO Conc. (mg/ml) | ASO Molarity (mM) | $Ca^{++}$ Molarity (mM) | $Mg^{++}$ Molarity (mM) | $MgCl_2$ suppl Molarity (mM) | $NaCl_2$ Molarity (mM) | Tonicity (mOsm/ml) | DivCat: ASO ratio |
|---|---|---|---|---|---|---|---|---|
| aCSF | 0 | 0 | 1.4 | 0.8 | 0 | 150 | **292** | **0** |
| dACSF+ ASO 100 mg | 100 | 14.1 | 1.4 | 0.8 | 0 | 42 | **286** | **0** |
| dACSF+7 mM total DivCat (Ca supp)+ASO 100 mg | 100 | 14.1 | 1.4 | 0.8 | 4.8 | 42 | **304** | **0.5:1** |
| dACSF+21 mM total DivCat (Ca supp)+ASO 100 mg | 100 | 14.1 | 1.4 | 0.8 | 18.8 | 0 | **260** | **1.5:1** |
| dACSF+42.2 mM total DivCat (Ca supp)+ASO 100 mg | 100 | 14.1 | 1.4 | 0.8 | 40 | 0 | **311** | **3:1** |

All groups were dosed at 30 mg/ 30µl. Abbreviations: DivCat= divalent cations; Ca supp= calcium supplemented; $MgCl_2$ suppl= magnesium supplemented; aCSF= standard artificial cerebrospinal fluid; dACSF= sodium deficient aCSF

**Table supplementary 7.** Formulation description of ASO A with or without magnesium supplementation in hypertonic or tonicity-controlled conditions (isotonic)

| Treatment group | ASO conc. (mg/ml) | ASO Molarity (mM) | $Ca^{++}$ Molarity (mM) | $Mg^{++}$ Molarity (mM) | $MgCl_2$ supp Molarity (mM) | $NaCl_2$ Molarity (mM) | Tonicity (mOsm/ml) | DivCat: ASO ratio |
|---|---|---|---|---|---|---|---|---|
| aCSF | 0 | 0 | 1.4 | 0.8 | 0 | 150 | **293** | 0 |
| aCSF+ 42.2 mM total DivCat (Mg supp) | 0 | 0 | 1.4 | 0.8 | 40 | 150 | **400** | 0 |
| aCSF+ ASO 100 mg | 100 | 14.1 | 1.4 | 0.8 | 0 | 150 | **492** | 0 |
| aCSF+ 42.2 mM total DivCat (Mg supp) + ASO 100 mg | 100 | 14.1 | 1.4 | 0.8 | 40 | 150 | **616** | 3:1 |
| dACSF+ 42.2 mM total DivCat (Mg supp) | 0 | 0 | 1.4 | 0.8 | 40 | 96 | **311** | 0 |
| dACSF+ ASO 100 mg | 100 | 14.1 | 1.4 | 0.8 | 0 | 42 | **285** | 0 |
| dACSF+ 42.2 mM total DivCat (Mg supp) + ASO 100 mg | 100 | 14.1 | 1.4 | 0.8 | 40 | 0 | **325** | 3:1 |

All groups were dosed at 30 mg/ 30µl. Abbreviations: DivCat= divalent cations; Mg supp= magnesium supplemented; aCSF= standard artificial cerebrospinal fluid; dACSF=sodium deficient artificial cerebrospinal fluid.

**Table supplementary 8.** Formulation description of ASO A with or without magnesium supplementation formulated in mACSF and aCSF in transgenic *ATXN3* mice

| Treatment group | ASO Conc. (mg/ml) | ASO Molarity (mM) | $Ca^{++}$ Molarity (mM) | $Mg^{++}$ Molarity (mM) | $MgCl_2$ supp Molarity (mM) | ASO dose (µg) |
|---|---|---|---|---|---|---|
| aCSF | 0 | 0 | 1.4 | 0.8 | 0 | 223 |
| aCSF+ 40 mM total $MgCl_2$ | 0 | 0 | 1.4 | 0.8 | 39.2 | 0 |
| aCSF+ ASO 22.3 mg | 22.3 | 3.14 | 1.4 | 0.8 | 0 | 223 |
| aCSF+ 40 mM total $MgCl_2$ + ASO 22.3 mg | 22.3 | 3.14 | 1.4 | 0.8 | 0 | 223 |
| mACSF+ 40 mM total $MgCl_2$ | 0 | 0 | 0 | 0 | 40 | 0 |
| mACSF+ ASO 22.3 mg | 22.3 | 3.14 | 0 | 0 | 0 | 223 |
| mACSF+ 40 mM total $MgCl_2$ + ASO 22.3 mg | 22.3 | 3.14 | 0 | 0 | 40 | 223 |

All mice received a dose volume of 10 µl. The dose is the calculated $ED_{80}$. Abbreviations: DivCat= divalent cations; $MgCl_2$= magnesium chloride; aCSF= standard artificial cerebrospinal fluid; mACSF=modified artificial cerebrospinal fluid.

**Table supplementary 9.** Formulation description of ASO D with or without $MgCl_2$ supplementation formulated in modified artificial cerebrospinal fluid (mACSF) in rats

| Treatment group | ASO Conc. (mg/ml) | ASO Molarity (mM) | $Ca^{++}$ Molarity (mM) | $Mg^{++}$ Molarity (mM) | $MgCl_2$ supp Molarity (mM) | ASO dose (mg) |
|---|---|---|---|---|---|---|
| mACSF | 0 | 0 | 0 | 0 | 0 | **0** |
| mACSF+ 40 mM total $MgCl_2$ | 0 | 0 | 0 | 0 | 40 | **0** |
| mACSF+ ASO 1 mg | 1 | 0.14 | 0 | 0 | 0 | **0.03** |
| mACSF+ ASO 3.3 mg | 3.3 | 0.46 | 0 | 0 | 0 | **0.1** |
| mACSF+ ASO 10 mg | 10 | 1.4 | 0 | 0 | 0 | **0.3** |
| mACSF+ ASO 33 mg | 33 | 4.6 | 0 | 0 | 0 | **1** |
| mACSF+ 40 mM total $MgCl_2$ + ASO 1 mg | 1 | 0.14 | 0 | 0 | 40 | **0.03** |
| mACSF+ 40 mM total $MgCl_2$ + ASO 3.3 mg | 3.3 | 0.46 | 0 | 0 | 40 | **0.1** |
| mACSF+ 40 mM total $MgCl_2$ + ASO 10 mg | 10 | 1.4 | 0 | 0 | 40 | **0.3** |
| mACSF+ 40 mM total $MgCl_2$ + ASO 33 mg | 33 | 4.6 | 0 | 0 | 40 | **1** |

All groups received a dose volume of 30µl. Abbreviations: DivCat= divalent cations; $MgCl_2$= magnesium chloride; mACSF= modified artificial cerebrospinal fluid.

**Table supplementary 10.** Formulation description of ASO A dosed at different concentrations and volumes with or without magnesium supplementation.

| Treatment group | ASO dose (mg) | Dose Volume (µl) | ASO Conc. (mg/ml) | ASO Molarity (mM) | $Ca^{++}$ Molarity (mM) | $Mg^{++}$ Molarity (mM) | $MgCl_2$ suppl Molarity (mM) | DivCat: ASO ratio |
|---|---|---|---|---|---|---|---|---|
| aCSF+ 42.2 mM total DivCat (Mg supp) | 0 | 30 | 0 | 0 | 1.4 | 0.8 | 40 | - |
| aCSF+ 42.2 mM total DivCat (Mg supp) | 0 | 100 | 0 | 0 | 1.4 | 0.8 | 40 | - |
| aCSF+ ASO 100 mg | 3 | 30 | 100 | 14.1 | 1.4 | 0.8 | 0 | 0.16:1 |
| aCSF+42.2 mM total DivCat (Mg supp)+ ASO 100 mg | 3 | 30 | 100 | 14.1 | 1.4 | 0.8 | 40 | 3:1 |
| aCSF+ ASO 30 mg | 3 | 100 | 30 | 4.22 | 1.4 | 0.8 | 0 | 0.52:1 |
| aCSF+ 12.7 mM total DivCat (Mg supp)+ ASO 30 mg | 3 | 100 | 30 | 4.22 | 1.4 | 0.8 | 10.5 | 3:1 |

Abbreviations: DivCat= divalent cations; Mg supp= magnesium supplemented; aCSF= standard artificial cerebrospinal fluid

**Table supplementary 11.** Detailed description of ASO A formulations used in Non-human primates' experiments.

| Treatment group NHP | ASO conc. (mg/ml) | ASO Molarity (mM) | Ca++ Molarity (mM) | Mg++ Molarity (mM) | $MgCl_2$ supp Molarity (mM) | Total DivCat Molarity (mM) | DivCat : ASO ratio | aA score (average) |
|---|---|---|---|---|---|---|---|---|
| aCSF+ ASO 70 mg | 70 | 9.9 | 1.4 | 0.8 | 0 | 2.2 | **0.2** | 3.25 |
| aCSF+ 9.2 mM total DivCat (Mg supp) + ASO 70 mg | 70 | 9.9 | 1.4 | 0.8 | 7 | 9.2 | **0.9** | 2 |
| aCSF+ 23.2 mM total DivCat (Mg supp) + ASO 70 mg | 70 | 9.9 | 1.4 | 0.8 | 21 | 23.2 | **2.3** | 0.87 |
| aCSF+ 23.2 mM total DivCat (Mg supp) + ASO 105 mg | 105 | 14.8 | 1.4 | 0.8 | 21 | 23.2 | **1.6** | 3.5 |
| aCSF+ 23.2 mM total DivCat (Mg supp) + ASO 140 mg | 140 | 19.7 | 1.4 | 0.8 | 21 | 23.2 | **1.2** | 4 |
| aCSF+ 33.2 mM total DivCat (Mg supp) + ASO 105 mg | 105 | 14.8 | 1.4 | 0.8 | 31 | 33.2 | **2.24** | 1.25 |
| aCSF+ 44.5 mM total DivCat (Mg supp) + ASO 140 mg | 140 | 19.7 | 1.4 | 0.8 | 42.3 | 44.5 | **2.25** | 2 |
| aCSF+ 58.3 mM total DivCat (Mg supp) + ASO 140 mg | 140 | 19.7 | 1.4 | 56.9 | 56.1 | 58.3 | **2.95** | 1 |

The volume injected in all groups = 1 ml. Abbreviations: DivCat=divalent cations; supp=supplemented; Aa score= acute activation score

**Table supplementary 12.** Primer sequences used in the study

| Target | Specie | Primers | |
|---|---|---|---|
| *Malat1* | Rat | Forward | CGTTAGACTTTTGTCACCTCACTTGA |
| | | Reverse | GGCAAGCGCTTATATGCAATC |
| | | Probe | TTTGCAGAGGCCTCATTTCATCCTTCA |
| *Gfap* | Rat | Forward | GAGAGAGATTCGCACTCAGTACGA |
| | | Reverse | GTCTGCAAACTTGGACCGATACCA |
| | | Probe | CAGTGGCCACCAGTAACATGCAAGAAAC |
| *Aif1* | Rat | Forward | AGGAGAAAAACAAAGAACACCAGAA |
| | | Reverse | CAATTAGGGCAACTCAGAAATAGCT |
| | | Probe | CCAACTGGTCCCCCAGCCAAGAX |
| *ATXN3* | Human | Forward | TGACACAGACATCAGGTACAAATC |
| | | Reverse | TGCTGCTGTTGCTGCTT |
| | | Probe | AGCTTCGGAAGAGACGAGAAGCCTA |
| *Gapdh* | Mouse | Forward | GGCAAATTCAACGGCACAGT |
| | | Reverse | GGGTCTCGCTCCTGGAAGAT |
| | | Probe | AAGGCCGAGAATGGGAAGCTTGTCATC |
| *Gapdh* | Rat | Forward | TGCTCCTCCCTGTTCTAGAGACA |
| | | Reverse | CACCGACCTTCACCATCTTGT |
| | | Probe | CCGCATCTTCTTGTGCAGTGCCAG |
